# A novel type of gel-like proteasome condensate induced by toxic protein aggregates

**DOI:** 10.1101/2025.09.15.676044

**Authors:** Liina Sirvio, Michael J. Morten, Matilda Burridge, Thomas Chua, Harry Whitwell, Yu Ye

## Abstract

Proteasomes reversibly form foci bodies in a liquid-liquid phase separation (LLPS)-dependent manner upon stress. We previously reported that internalized protein aggregates were targeted by proteasome-dense foci^1^, and proposed that such transient aggregate-associated droplets (TAADs) may facilitate aggregate removal^2^. Here we use quantitative imaging to show that TAADs represent a novel type of gel-like proteasome condensate. TAADs are irregular in shape and slow to disperse, sequestering proteasomes in agreement with our observation of confined diffusion^3^. We demonstrate that TAADs co-localize with cytosolic alpha-synuclein aggregates to facilitate their clearance. Inhibition of proteasome- or ubiquitination activity abolishes this aggregate clearance. We identify RAD23B necessary for TAAD formation, amid other co-localizing chaperones and (co-)proteins of the ubiquitin-proteasomes system. TAAD formation is associated with higher proteasomal substrate turnover whilst retaining overall catalytic efficiency, suggestive of altered degradation mechanisms upon aggregate engagement. Proteomics analysis reveals impact on key mitochondrial-associated processes even after TAAD-aggregate disengagement. Similar TAAD-aggregate co-localizations are found in iPSC-differentiated neurons and in disease-relevant samples, with no detection of compromised proteasome activity. Together, our results indicate a model where TAADs concentrate local proteasome activity, which facilitates aggregate clearance in healthy ageing cells. Potentially, should pathological aggregates persist, TAADs may remain engaged and conceivably sequester proteasomes from physiological activities, thus contributing to neurodegenerative disorders.

## Introduction

Selective protein degradation by the ubiquitin-proteasome system (UPS) is an important cellular mechanism that modulates vital processes such as transcriptional regulation, DNA repair, inflammatory-, oxidative stress- and unfolded protein responses^4–6^. The localization of proteasomes is highly dynamic, reflecting a regulated distribution of cellular degradation activity that may change to adapt to stress conditions^7–11^. A process termed liquid-liquid phase separation (LLPS) has more recently been reported to transiently alter localization through condensation of proteasomal particles, which are normally distributed throughout the cell, into foci bodies^12^. Behaving like liquid droplets with increased degradation activity, these LLPS-foci have been known to be induced by osmotic stress^13,14^, starvation^15^, and cell senescence^16^. Whether foci induced by distinct types of stress share similar protein compositions and functions remains unresolved^17^.

Locally concentrating degradation activity may enable clearance of protein aggregates^12,18^, thus reducing proteotoxic stress. We recently proposed that proteasomes may concentrate and condense together with chaperones, UPS enzymes as well as co-proteins such as proteasome shuttling factors, in putative transient aggregate-associated droplets (TAADs) that can facilitate aggregate clearance^2^. In support of this proposition, we reported that internalized alpha-synuclein (aS) aggregates, which are hallmarks of Parkinson’s disease (PD), co-localized with proteasome-intense foci and led to gradually diminished aggregate size^1^, in alignment with our expected TAAD function. Using multiple biophysical techniques, we also recently reported the molecular mechanisms of aggregate-induced TAAD formation and attributed this to confined proteasome diffusion^3^. It remains unclear whether such TAADs resemble LLPS condensates and what impact their formation has on canonical proteostasis events and signaling processes regulated by the UPS.

Here, we report that aS aggregates induce a novel type of proteasome condensate, which did not readily exchange with the surrounding cytoplasm nor dissolve in aliphatic alcohol. In agreement with our previous proposition of TAADs^2^, these condensates co-localized with internalized aggregates within 24 hrs and disappeared after 48 hrs, when aggregate load reduced. Both TAAD formation and aggregate removal were sensitive to ubiquitination and proteasome inhibition, assisted by key proteasomal shuttling factors. Reconstructed 2D and 3D super-resolution images by single-molecule localization microscopy (SMLM) showed that TAADs are irregularly shaped, unlike the largely spherical LLPS-foci. TAAD formation was further validated in iPSC-derived cortical-like neurons and found in neurons from Parkinson’s disease (PD) donor tissues. Critically, we performed full Michaelis-Menten kinetics analysis and did not detect reduced proteasome activity in model cells and donor tissues, while proteomics analysis revealed lasting changes to proteins in mitochondrial-associated processes. Together, our data indicate that TAADs are a previously uncharacterized type of proteasome condensate that facilitates aggregate removal. We propose a model where internalization of aggregates induces formation of TAADs, which temporarily sequester proteasomes from partaking in cellular proteostasis, whilst prolonged TAAD formation will compromise proteostasis and physiological processes, thus contributing to pathology.

## Results

### Stress induces proteasome re-distribution

We investigated proteasome dynamics in CRISPR-edited HEK293A cells expressing either PSMD14-eGFP or PSMB2-eGFP, subunits of the proteasome 19S regulatory-(RP) or 20S core particle (CP), respectively^1^. Highly inclined and laminated optical sheet (HILO) imaging was performed on cells grown on coverslips^1^ (see **Materials and Methods**). RP and CP were found distributed throughout the cell with high density in the nucleus (**Fig.1A** and **SFig.1A**). We have described the distribution and transport mechanisms of proteasomes elsewhere^3^, confirming that most proteasome particles are moving and functioning as holoenzymes in cells^19,20^. Inducing osmotic shock or hypertonic stress with established methods^12,13^ using sucrose or CaCl_2_, respectively, resulted in the formation of proteasome LLPS-foci within 5 min (**Fig.1B-C** and **SFig.1B-C**). To investigate proteasomes as TAADs, we quantified cells at 3, 24 and 48 hrs post-incubation with aggregates assembled from recombinant aS^1^ (**Fig.1D** and **SFig.1D-E**. See also **Materials and Methods**). TAADs were detected in the nucleoplasm within 3 hrs, and co-localized with cytosolic aggregates at 24 hrs, eventually disappearing at 48 hrs (**Fig.1E-F** and **SFig.1F**). Compared to LLPS-foci, proteotoxic stress-induced TAADs were fewer in number but much larger, as quantified by HILO imaging (**Fig.1G**). Co-localization was only observed at 24 hrs (**Fig.1F**) followed by concomitant decrease in the number of aggregates (**Fig.1H** and **Sig.1G**) and of TAADs (**Fig.1I** and **SFig.1H**) at 48 hrs, suggesting that interaction with TAADs facilitated aggregate clearance.

**Figure 1.**
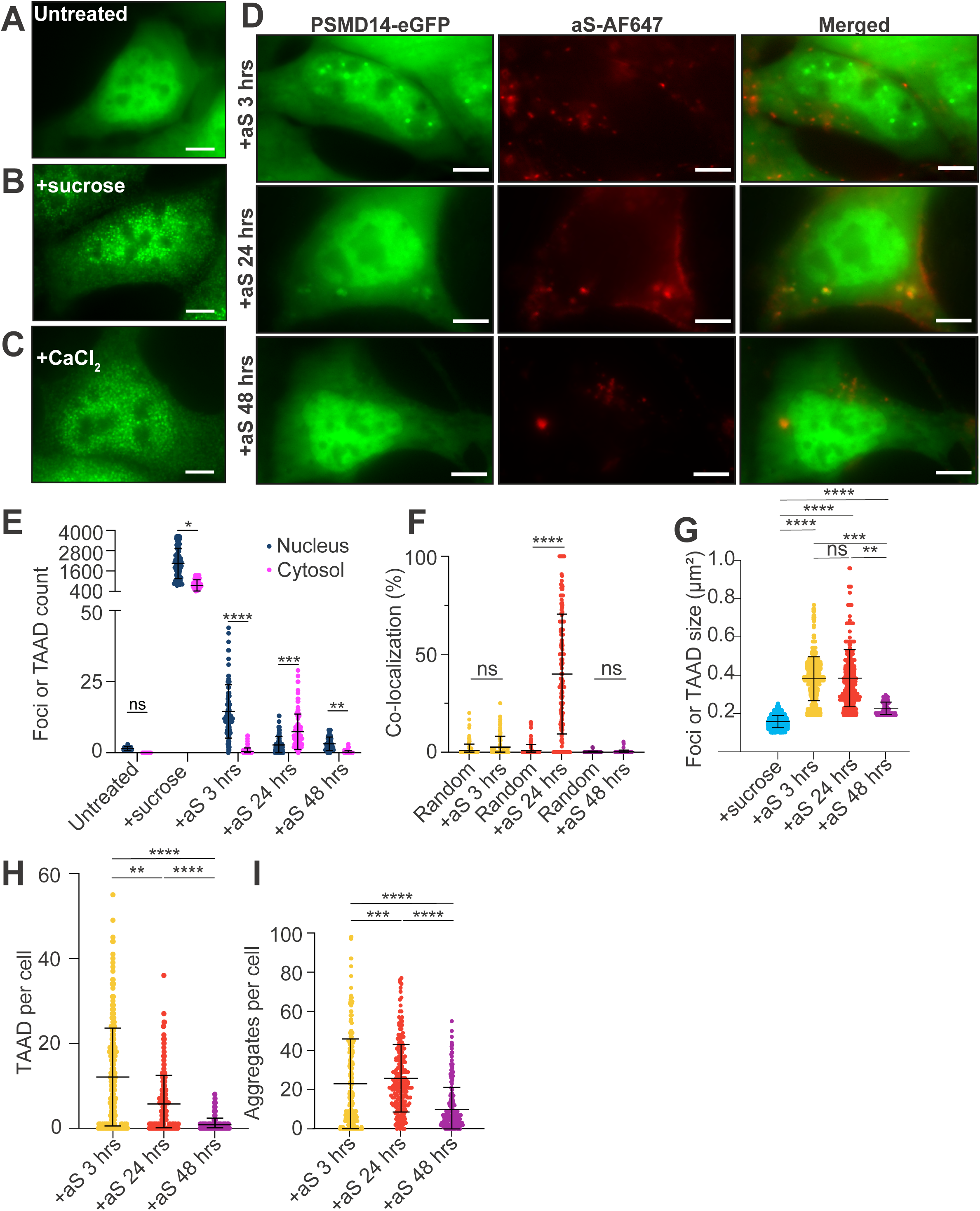
TAADs co-localize with internalized aS aggregates. **(A)** Live-cell imaging of resting or **(B)** 5 min post incubation with 200 mM sucrose or (**C**) 150 mM CaCl_2_ of CRISPR-engineered PSMD14-eGFP cells (see **Materials and Methods**). Similar results were observed for PSMB2-eGFP cells in **SFigure 1A-C**. **(D)** PSMD14-eGFP cells were incubated for 3, 24 or 48 hrs with AF647-labeled aS aggregates (red) at 1 μM. Images are representative of three independent experiments (n>200 cells for each condition, same below unless otherwise stated). Scale bars represent 5 μm. See **SFigure 1D** for similar observations in PSMB2-eGFP cells. Final concentrations are given for sucrose and aS and used in subsequent experiments, unless otherwise stated. (**E**) Quantification of the number of foci contained in the nucleus and cytosol for untreated cells (nuclear: 1.5 ± 0.7, cytosolic: 0.0 ± 0.0, n=12 cells), cells incubated with sucrose (nuclear: 2061 ± 898, cytosolic: 762 ± 332, n=84 cells), with aS aggregates for 3 hrs (nuclear: 14.6 ± 9.4, cytosolic: 0.5 ± 1.2, n=88 cells), 24 hrs (nuclear: 2.7 ± 3.0, cytosolic: 7.4 ± 6.2, n=103 cells) or 48 hrs (nuclear: 3.2 ± 2.4, cytosolic: 0.2 ± 0.7, n=45 cells). Data are mean ± SD and were analyzed by multiple t-test. (**F**) Co-localization of TAADs with aggregates compared with chance co-localization (random) after 3 hrs (2.6 ± 5.5, n=149 cells; random: 1.0 ± 3.1), 24 hrs (39.8 ± 30.5, n=172 cells; random 0.9 ± 3.0) or 48 hrs (0.2 ± 0.8, n=114 cells; random: 0.1 ± 0.5). **(G)** Comparison between the areas of proteasome foci bodies induced by different stimuli (sucrose = 0.16 ± 0.03 µm^2^, foci=17,599, n=84 cells; TAADs at 3 hrs: 0.38 ± 0.11 µm^2^, TAAD=1588, n=262 cells; at 24 hrs: 0.38 ± 0.15 µm^2^, TAAD=1157, n=209 cells; 48 hrs: 0.23 ± 0.03 µm^2^, TAAD=124, n=229 cells). (**H**) Quantification of TAAD number per cell at 3 hrs (12.0 ± 11.5, n=261 cells), 24 hrs (5.7 ± 6.9, n=209 cells) or 48 hrs (0.7 ± 1.6, n=229 cells) incubation with aS aggregates. (**I)** Quantification of the number of aggregates per cell following 3 hrs (23 ± 22), 24 hrs (25 ± 17) or 48 hrs (10 ± 11) incubation as in **H**. At least three biological repeats were performed for each condition and combined for presentation. All data are mean ± SD by t-test.

These observations suggest that TAAD formation is initially observed in the nucleus where proteasome concentration is highest^3^, and eventually emerge in the cytosol to co-localize with aggregates, leading to their clearance and re-dispersal of proteasomes at 48 hrs. Monomeric aS or aS aggregation buffer alone did not induce TAADs (**SFig.2**), confirming that the formation of TAADs was specific to the presence of aS aggregates.

**Figure 2.**
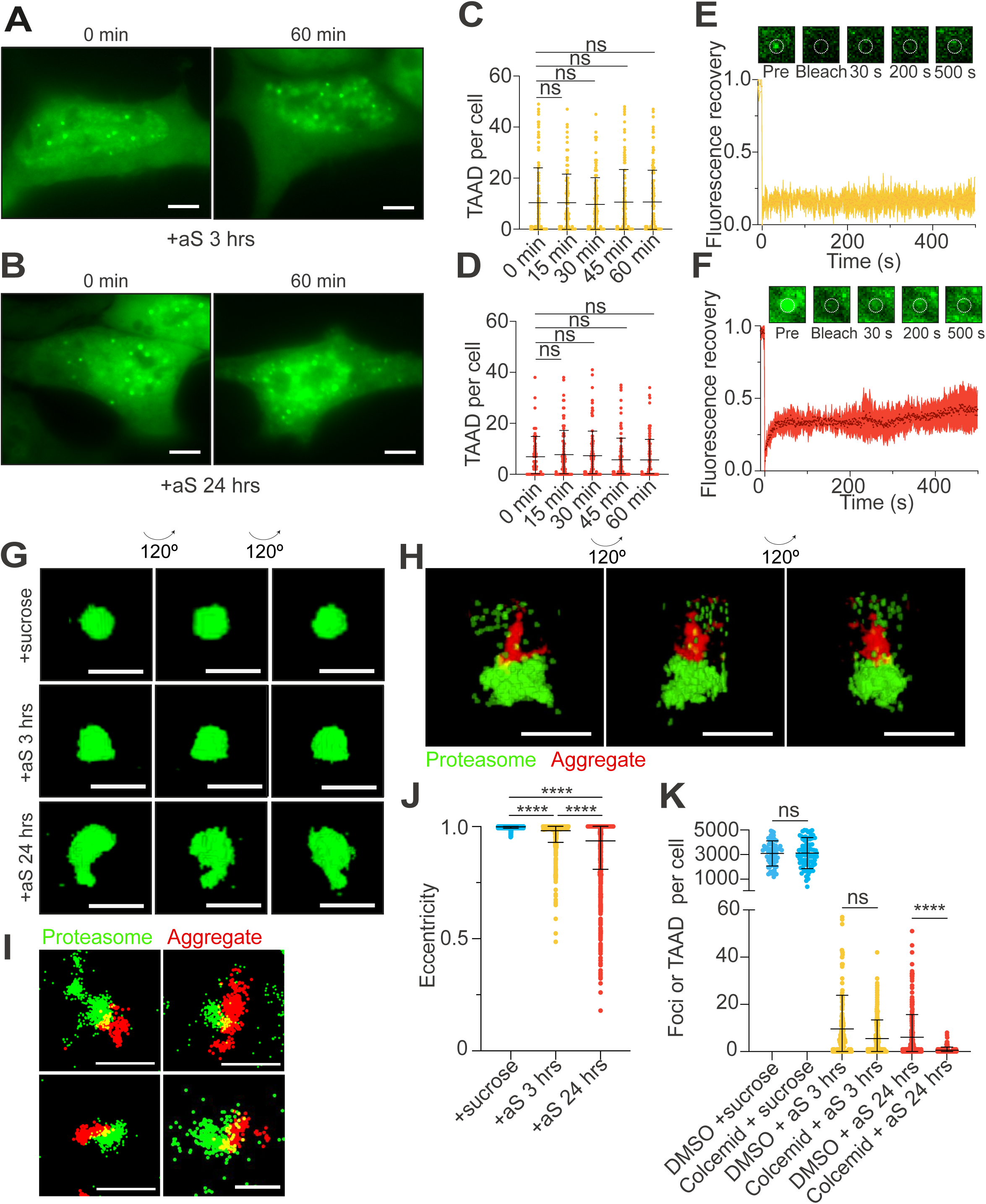
TAADs are gel-like condensates and distinct to LLPS-foci. (**A-B**) Representative images of PSMD14–eGFP cells incubated with aS aggregates at 1 μM for **(A)** 3 hrs or **(B)** 24 hrs, and subsequently treated with 5% 1,6-hexanediol (1,6-HD) for 60 min. Scale bars represent 5 μm. (**C-D**) Number of TAADs observed per cell were quantified at multiple timepoints post for 1,6-HD treatment for TAADs at 3 hrs (0 min: n=122 cells; 15 min: n=110 cells; 30 min: n=118 cells; 45 min: n=115 cells; 60 min: n=124) and 24 hrs (0 min: n=96 cells; 15 min: n=93 cells; 30 min: n=108 cells; 45 min: n=111 cells; 60 min: n=94 cells). Data are presented as mean ± SD by t-test and from at least three independent repeats. **(E-F)** Quantification of FRAP experiments fitted to single-exponential functions for TAADs at 3 (*top*) and 24 hrs (*bottom*). Results are mean ± SD of three independent experiments. See **S**Figure 3 for 1,6-HD and FRAP experiments performed on LLPS-foci, the positive control. **(G)** Reconstructions in 3D of sucrose- and aggregate-induced proteasome condensates, created from z-stacks of images taken 100 nm apart. Scale bars represent 1 μm. See also **SVideos 1-3**. (**H**) SMLM in 3D and (**I**) in 2D of live PSMD14-mEos and PSMB2-mEos cell lines, respectively, incubated with aS aggregates for 24 hrs. Scale bars represent 0.5 and 1 μm for 3D and 2D images, respectively. See also **SVideo 4**. (**J**) Mean ± SD eccentricity scores on LLPS-foci (eccentricity = 0.998 ± 0.006, foci=17,002, n=84 cells) and TAADs (3 hrs: eccentricity = 0.981 ± 0.052, TAAD=1934, n=262 cells; 24 hrs: eccentricity = 0.936 ± 0.127, TAAD=1490, n=209 cells) were analyzed by t-test. (**K**) PSMD14-eGFP cells were incubated with (*from left*): 2 μM colcemid or DMSO vehicle control for 1 hr prior to treatment with sucrose (DMSO+sucrose: 3096 ± 1026, n=77 cells; colcemid+sucrose: 2462 ± 1227, n=89 cells); or incubated with 2 μM colcemid or DMSO and aS aggregates for 3 hrs prior to imaging (DMSO+aS: 9.2 ± 13.3, n=175 cells; colcemid+aS: 5.5 ± 7.9, n=242 cells); or with 0.25 μM colcemid and aS aggregates for 24 hrs prior to imaging (DMSO+aS: 5.1 ± 7.8, n=145 cells; colcemid+aS: 0.58 ± 1.4, n=114 cells). Data are presented as mean ± SD by t-test.

### TAADs demonstrate gel-like properties

We speculated whether TAADs would also be LLPS-dependent entities and tested this with 1,6-hexanediol, which is canonically used to disrupt hydrophobic interactions. The number and size of TAADs formed after 3 and 24 hrs aggregate incubation remained up to 60 min after treatment (**Fig.2A-D**), whereas sucrose-induced LLPS-foci expectedly dissolved^12^ within 5 min by 1,6-hexanediol (**SFig.3A**). This suggests that interactions that maintain TAAD formation are distinct to those in LLPS-foci. In support of this, fluorescence recovery after photobleaching (FRAP) experiments showed rapid recovery in LLPS-foci comparable to that of the cytosolic background (**SFig.3B-C**), while little to no recovery was observed for TAADs (**Fig.2E-F**). These experiments suggest that TAADs likely represent gel-rather than liquid-like condensates, and diffusional exchange of proteins across the phase boundary is slow. Since TAADs at 3 and 24 hrs both show similar properties, we conclude that their gel-like nature is independent from interacting with aggregates.

**Figure 3.**
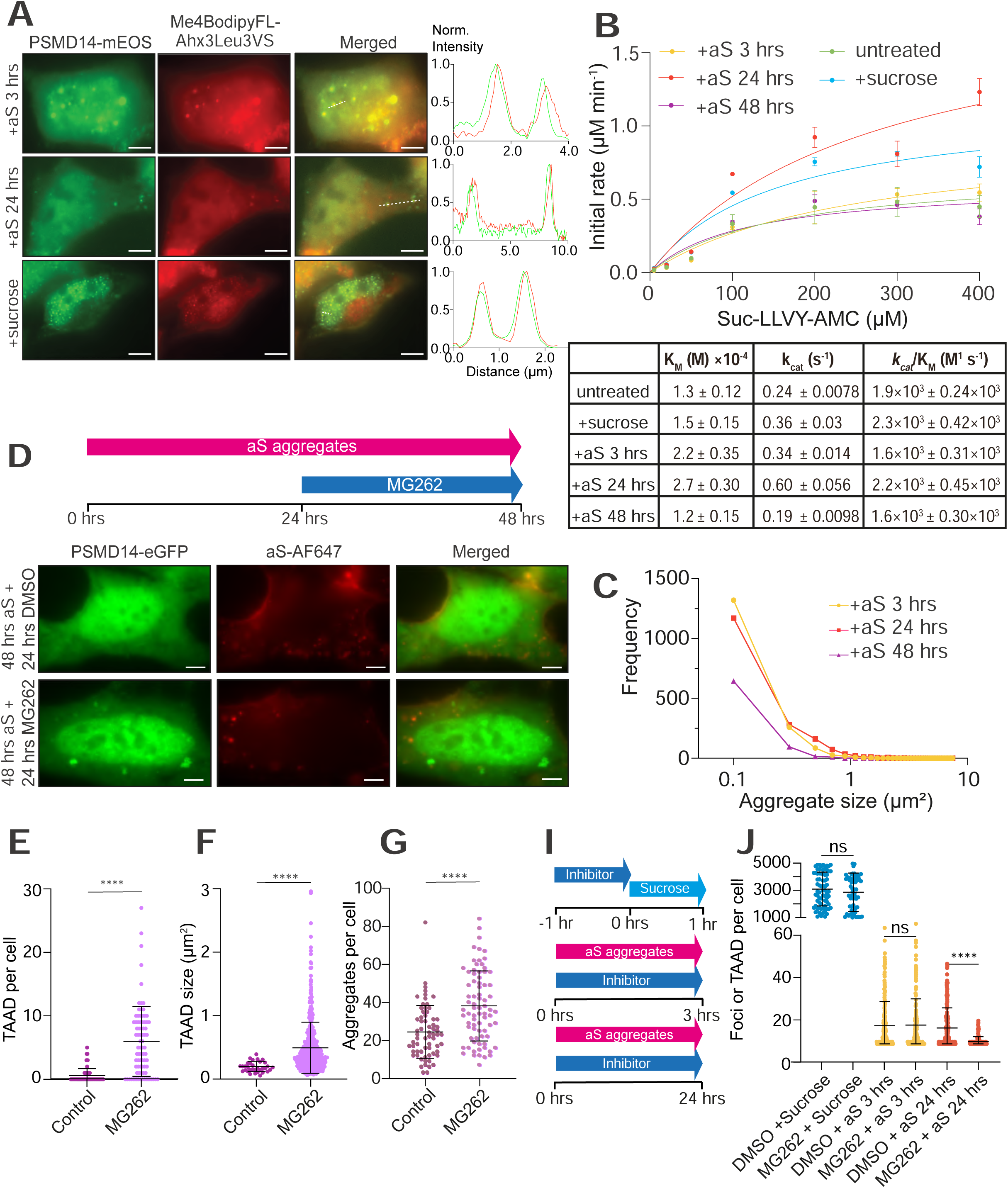
TAADs are active degradation centers for aggregates. **(A)** Proteasome degradation measured by activity probe Me4BodipyFL-Ahx3Leu3VS (red) stained for 1 hr in live PSMD14-eGFP cells (green) pre-incubated with aS aggregates for 3 (*top*) or 24 hrs (*middle*), or with sucrose (*bottom*). Images are representative of three independent experiments. Scale bars = 5 μm. Line profiling of representative sections of cells, indicated by white dashed lines, are shown on the right and color-coded accordingly. **(B)** Michalis-Menten functions were fitted to initial reaction rates measured over a concentration range of a model substrate, LLVY-AMC, incubated with a constant lysate amount of known proteasome concentration (see **S**Figure 4B and **Materials and Methods**). Lysates from control cells, and from cells incubated with sucrose or aS aggregates for 3, 24 or 48 hrs were measured separately. Catalytic efficiencies were calculated from the Michalis constant, K_M_, and the turnover number, *k_cat_*, for each condition. Error bars indicate SEM from three independent repeats. **(C)** Change in the level of internalized aggregates binned at different sizes (given in image area) at 3, 24, or 48 hrs post incubation. Aggregate numbers from cells in **Figure 1I** were combined for presentation. (**D-G**) PSMD14-eGFP cells were incubated with 1 μM aS aggregates for 48 hrs and treated with 0.25 μM MG262 or DMSO vehicle control for an additional 24 hrs to inhibit proteasome activity, as depicted in **(D**). Quantifications of (**E**) number of foci per cell (control: 0.5 ± 1.11, n=66 cells; MG262: 5.93 ± 5.55, n=80 cells), (**F**) TAAD area (control: 0.19 ± 0.014, TAAD=33, n=66 cells; MG262: 0.485 ± 0.018, TAAD=484; n=80 cells) and (**G**) number of aggregates per cell (control: 24.30 ± 1.71, n =66 cells; MG262: 38.03 ± 2.10, n=77 cells) are shown in mean ± SD by Mann-Whitney test. **(I)** Schematic description of 50 µM MG262 treatment in PSMD14-eGFP cells for 1 hr prior to sucrose treatment to test effect on LLPS-foci formation, or concurrently with aS aggregate incubation for 3 or 24 hrs to test effect on TAAD formation. **(J)** Quantification of treatment with MG262 or DMSO vehicle control performed as in **I** to examine LLPS-foci formation (DMSO+sucrose: 3107 ± 1251, n=73 cells; MG262+sucrose: 2870 ± 1420, n=77 cells), TAAD at 3 hrs (DMSO+aS: 8.6 ± 11.5, n=189 cells; MG262+aS: 8.9 ± 12.5, n=169 cells) and TAAD at 24 hrs (DMSO+aS: 7.5 ± 9.6, n=167 cells; MG262+aS: 1.1 ± 2.3, n=189 cells). Data are mean ± SD and analyzed by t-test.

The shapes of TAADs and LLPS-foci were subsequently reconstructed from *in situ* imaging. Compared to the predominantly spherical LLPS-foci (**Fig.2G**), all TAADs examined were irregularly shaped and of individually distinct appearances that may complement the shapes of respective aggregates with which they are interacting (**Fig.2H-I**). In agreement, the eccentricity score for LLPS-foci was higher than TAADs, confirming their morphological differences (**Fig.2J**). Using SMLM, we observed a clear interface in all examined interactions between the aggregate and the TAADs at 24 hrs (**Fig.2I**), and visualized this interaction in 3D (**Fig.2H**). These observations demonstrate direct TAAD-aggregate engagement, likely facilitating aggregate clearance.

The clear interface may result from TAADs that have pre-formed before being transported to target aggregates, in line with TAADs at 3 hrs that showed no co-localization (**Fig.1D** and **F**). We therefore examined involvement of cytoskeletal transport and depolymerized microtubules by adding colcemid to cells pre-incubated with aggregates, which showed that TAAD formation was indeed impeded (**Fig.2K**). This is also consistent with the gradually shifting ratio of nuclear to cytoplasmic TAADs over time (**Fig.1E**). It is tempting to suggest that TAADs may have moved from their formation in the nucleus at 3 hrs to their appearance in the cytosol at 24 hrs, at which point co-localization with aggregates was highest (**Fig.1E-F**), though further direct evidence would be required to demonstrate such transport. Together, our observations demonstrate that TAADs represent a novel type of gel-like proteasome condensate that target aggregates and distinct to previously reported proteasomal condensates.

### TAADs serve as degradation centers

Whether proteasomes can degrade or are inhibited by specific aS aggregate forms has been debated^21–23^. We therefore compared proteasome activities of TAADs in live cells using a fluorescence reporter probe (Me4BodipyFL-Ahx3Leu3VS)^24^, which showed dispersed proteasomal activity that correlated with proteasome distribution in resting cells (**SFig.4A**). Fluorescence intensities from the reporter probe increased for TAADs in aggregate-incubated cells at 3 and 24 hrs, as did intensities detected in LLPS-foci, the positive control^12^ (**Fig.3A**), suggesting concentrated degradation activity rather than proteasome inhibition. To validate this observation and quantify changes in degradation activity upon aggregate internalization, we obtained lysates from cells incubated under different conditions and measured the rates of reaction using LLVY-AMC, a fluorescent model substrate. This enabled us to determine Michaelis-Menten functions of proteasome kinetics for each sample, from which we calculated catalytic efficiencies (*k_cat_*/K_M_) (**Fig.3B** and **SFig.4B**). Our data showed that K_M_ gradually increased with aggregate incubation from resting to 3 and 24 hrs, and returned to base level at 48 hrs. Meanwhile *k_cat_* remained at similar levels in resting cells or cells 3 or 48 hrs post-incubation, and saw a meaningful increase at 24 hrs, when TAAD-aggregate co-localization was highest in **Fig.1F**.

**Figure 4.**
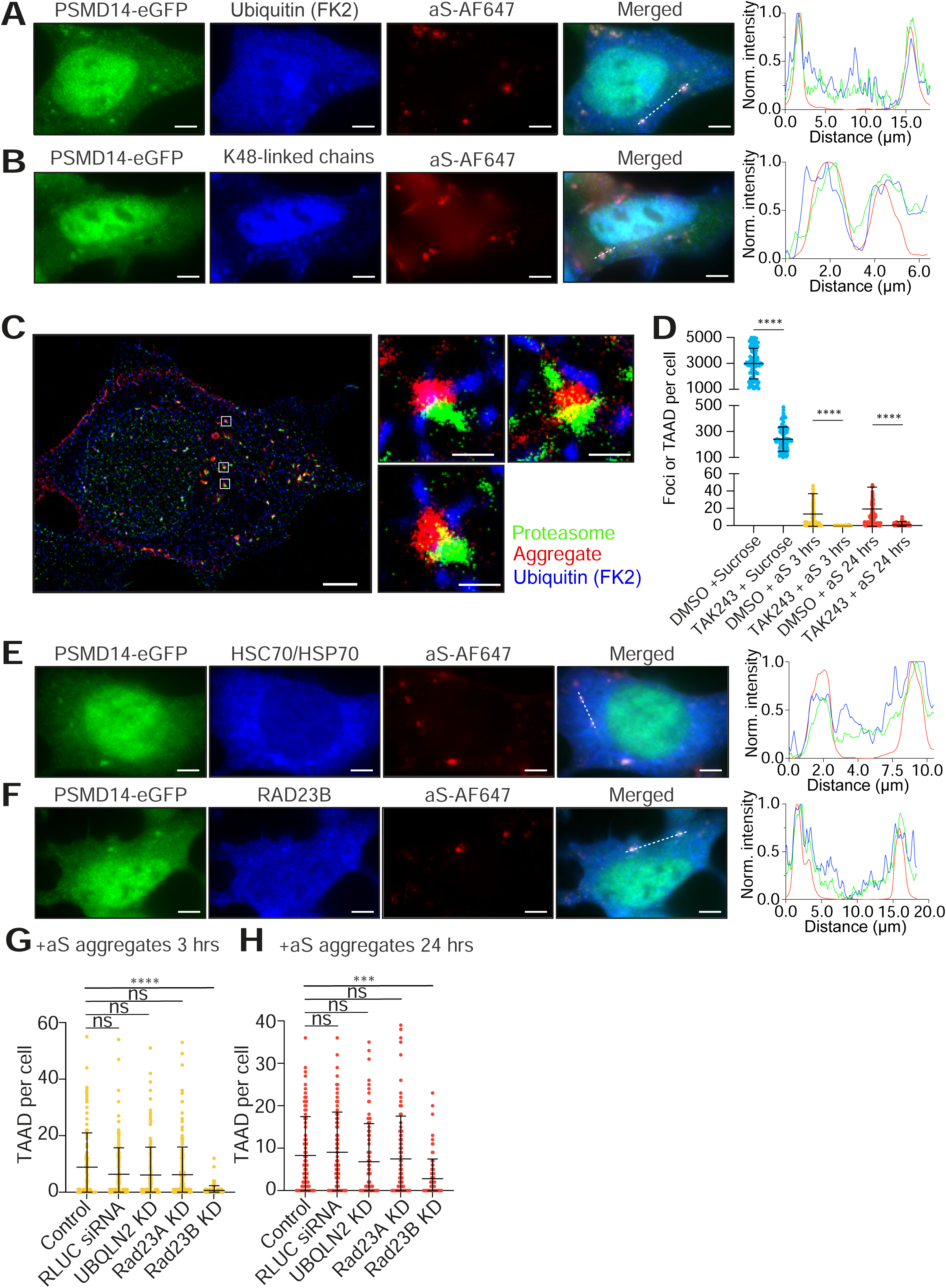
TAADs contain chaperones, shuttling factors and UPS enzymes. PSMD14-eGFP (green) cells were incubated with aS aggregates (red) for 24 hrs and fixed as described in **Materials and Methods** prior to immunostaining with respective antibodies (in blue) for **(A)** ubiquitin (FK2) or **(B)** Lys48-linked ubiquitin chain. Scale bars = 5 µm. Line profiles of respective cell sections as indicated by white dashed lines are color-coded accordingly. Images are representative of at least three biological repeats. **(C)** Representative SMLM image of a typical PSMD14-mEos cell (green) incubated with AF647-labeled aS aggregate (red) and stained with antibody against ubiquitin (FK2, blue). Scale bars represent 5 μm or 1 μm (zoomed-in). **(D)** TAK243 treatment at 1 µM or DMSO vehicle control followed the same scheme as in **Fig.3I** to examine effect on LLPS-foci (DMSO+sucrose: 3003 ± 1186, n=94 cells; TAK243+sucrose: 240.4 ± 93.3, n=99 cells), TAADs after 3 hrs (DMSO+aS: 6.2 ± 10.05, n=230 cells; TAK243+ aS: 0.04 ± 0.07, n=224 cells) or TAADs after 24 hrs (DMSO+aS: 8.6 ± 10.9, n=138 cells; TAK243+aS: 0.7 ± 1.6, n=127 cells). Data are mean ± SD and analyzed by t-test. Immunostaining against **(E)** HSC70/HSP70 and **(F)** RAD23B performed and presented as in **A**. (**G-H**) Quantification of TAADs following knock-down (KD) in PSMD14-eGFP cells 48 hrs post-transfection with RLUC siRNA, siRNA targeting UBQLN2, RAD23A and RAD23B, or no siRNA control were incubated with aS aggregates for 3 hrs (control: 8.7 ± 12.1, n=104 cells; RLUC siRNA: 6.4 ± 9.3, n=146 cells; UBQLN2: 6.1 ± 9.9, n=134 cells; RAD23A: 6.3 ± 10.3, n=227 cells; RAD23B: 0.7 ± 1.6, n=138 cells) or 24 hrs (control: 8.3 ± 9.2, n=80 cells; RLUC siRNA: 9.1 ± 9.5, n=71 cells; UBQLN2: 6.9 ± 9.0, n=85 cells; RAD23A: 7.6 ± 10.2, n=97 cells; RAD23B: 2.9 ± 4.6, n=83 cells). Number of TAADs were counted from at least three independent repeats and plotted as mean ± SD. See **SFigure 7** for siRNA KD validation.

Our interpretation of the data is that although the affinity of proteasomes for the model substrate (LLVY-AMC) reduced upon TAAD-aggregate co-localization, perhaps to prioritize aggregate engagement, the rate of substrate turnover is increased to maintain catalytic efficiency in the cell, thus compensating for proteasomes that have engaged aggregates. This is consistent with proteasomes that accumulated with neurotoxic poly-GlyAla aggregates in neurons were observed in the catalytically active conformation^25^. Indeed, catalytic efficiencies for sucrose-stressed cells were also within a similar range to proteotoxic-stressed cells and control cells (**Fig.3B**). Careful examination of fluorescence intensities of TAADs to determine ratio of RP:CP revealed that while the number of both RP and CP increased from 3 to 24 hrs, CP was more than twice abundant as RPs (**SFig.4C-E**). As proteolytic activities reside in the CP while RPs facilitate ubiquitin recognition and substrate binding, the altered CP:RP ratio potentially contributed to the higher substrate turnover and reduced affinity, respectively. Together, these results suggest a carefully tuned cellular mechanism to maintain proteostasis processes under resting and stress conditions.

Examining the substrates, we found that mostly small aggregates (<500 nm) were internalized within 3 hrs incubation, in alignment with our previous findings^1^. Their level remained between 3-24 hrs, suggesting no overall aggregate clearance during this time window (**Fig.3C**). A striking decrease in the number of aggregates across all sizes followed between the 24-48 hrs timespan (**Fig.3C**), coinciding with TAAD-aggregate co-localization timepoint in **Fig.1**. These results support that proteasomes target internalized aggregates and gradually process them into smaller fragments for further degradation^1,22,23^ rather than targeting particular aggregate species.

This proteasome-dependent aggregate degradation was validated by treating cells already pre-incubated with aggregates with vehicle control or MG262 at low concentration (see chart, **Fig.3D**). Proteolytic-specific inhibition with MG262 quantitatively prevented aggregate clearance compared to vehicle control (**Fig.3E**). Instead, TAADs increased both in abundance and size when examined 48 hrs after aggregate incubation (**Fig.3E-G**). MG262 is not involved in facilitating or retaining TAADs, as simultaneous MG262 treatment and aggregate incubation abolished TAAD formation (**Fig.3I-J**). These results indicate that observed aggregate clearance is facilitated by proteasomes and TAAD formation, in agreement with our previous biochemical^22,23^ and cellular^1^ findings that proteasomes target and reduce aggregate size. Together, our observations exemplify the role of proteasomes in aggregate degradation, and the possibility that once formed, TAADs can re-disperse only after aggregate clearance. This may imply that persistent aggregate engagement results in augmented TAAD formation.

### UPS enzymes, chaperones and shuttling factors contribute to TAAD activity

We next sought evidence for whether ubiquitination is involved in mediating aggregate degradation and performed immunofluorescence staining using an optimized fixation protocol that partially retained TAADs at 24 hrs (see **SFig.5** and **Materials and Methods**). Pan-ubiquitin antibody (P4D1, **SFig.6A**) and an antibody specific for conjugated ubiquitin (FK2, **Fig.4A**) both co-localized with TAAD-aggregate complexes, suggesting presence of protein ubiquitination. To test for polyubiquitination, we stained against Lys48- and Lys63-linked chains, the two most common linkages, and found their presence in the TAADs (**Fig.4B** and **SFig.6B**). The ubiquitination may have been catalyzed, at least in part, by co-localization with E2 enzymes such as UBE2D/UbcH5, which may assemble both chain types depending on the E3 ligase^26^ (**SFig.6C**). STUB1/CHIP is known to ubiquitinate aggregates^27^ and efficiently target misfolded proteins such as tau and aS^28,29^, was absent from TAADs (**SFig.6D**), indicating that other ligases may be involved. Alternatively, aggresome-associated free ubiquitin chains have also been shown to recruit proteasomes^30^, providing another possibility for observed Ub chains. To test this, we performed multi-color SMLM on TAADs at 24 hrs and indeed detected conjugated ubiquitin on aggregates (**Fig.4C**). Treating cells with TAK243, the inhibitor against E1 enzymes, demonstrated that that ubiquitination is necessary for TAAD formation (**Fig.4D**) and recruitment. These observations together suggest a model where ubiquitination of aggregates lead to recruitment of TAADs. This is also supported by enrichment of ubiquitin- and degradation-related proteins in the proteomics (see below **Fig.5**).

**Figure 5.**
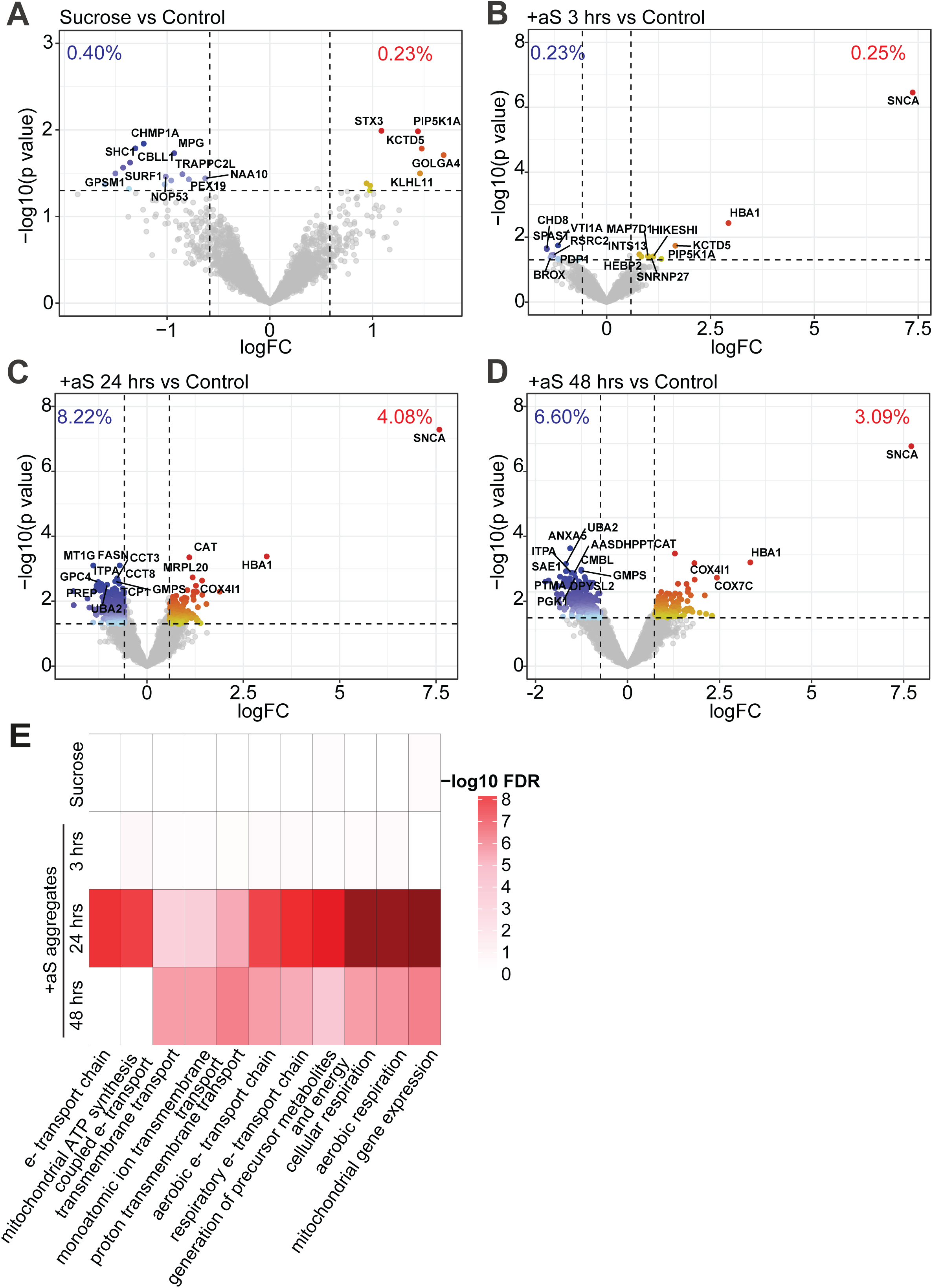
Aggregate internalization causes lasting impact on distinct cellular functions. **(A-D)** Analysis of proteomic changes in the pellet after incubation with **(A)** sucrose for 45 min, or with aS aggregates for **(B)** 3 hrs, **(C)** 24 hrs or **(D)** 48 hrs in PSMD14-eGFP cells (n=3 biological replicates). Results are presented as volcano plots with log_2_-fold change versus – log_10_ p-value for each condition compared with control. Proteins meeting the significance threshold (|log_2_FC| > log_2_(1.5), p-value < 0.05) are colored on a gradient scale according to spectral-count adjusted p-value (upregulated: yellow to red; downregulated: light to dark blue). Non-significant proteins are displayed in grey. Percentage of significantly up- and down-regulated proteins is indicated on each plot, and the top 15 candidates are labeled by gene name. See also **Supplementary Table 1**. **(E)** Heatmap of gene ontology (GO Biological Processes) over-representation of up-regulated proteins across all treatments. Displayed as top 10 terms from each treatment, curated to reduce term redundancy. Colored according to −log_10_(FDR) (Benjamini–Hochberg adjusted p-values).

**Figure 6.**
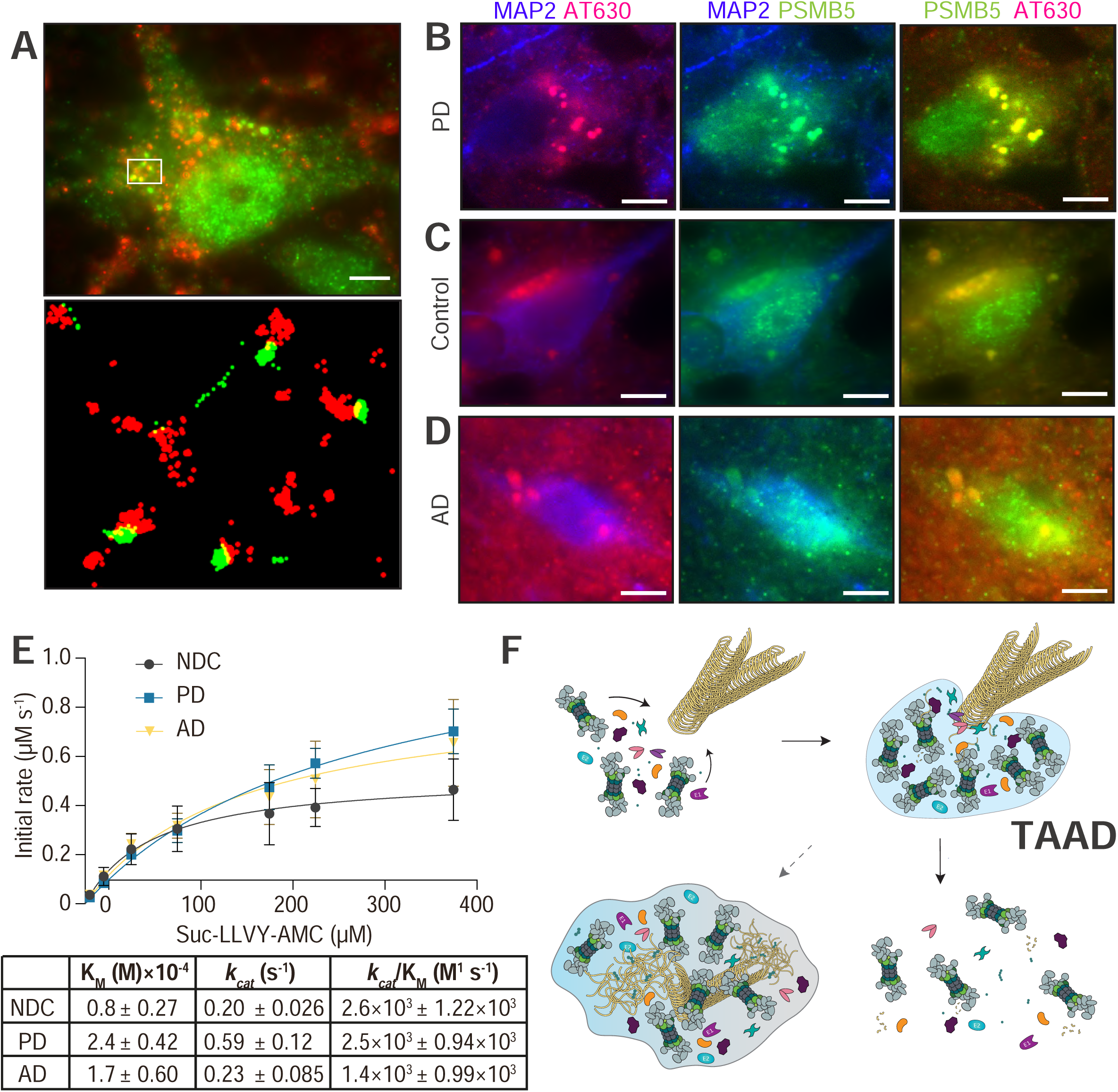
TAAD formation in model pathological models. (**A**) Neurons at >100 DIV differentiated from a human iPSC line carrying an *SNCA* triplication mutation subjected to aggregate incubation as in **Figure 1D** for 24 hrs. Proteasomes (anti-PMSB5, green) co-localized with AF647-labeled aS aggregates (red), with reconstructed SMLM image of the boxed region shown zoomed-in below. Scale bar = 5 µm. See also **SFigure 10A** individual channels. **(B-D)** Representative images of neurons in frontal cortical brain tissues donated by a **(B)** PD **(C)** non-disease control or **(D)** AD donor. Immunostaining was performed against a neuronal (MAP2, blue) and a proteasome marker (PSMB5, green). Aggregates were stained in situ using Amytracker-630 (AT630, red), an imaging approach we previously reported and worked well to detect co-localization between TAADs and internalized pathological aggregates^1^. Scale bars = 5 µm. The original color intensity in **C** was weak and increased tenfold for presentation. (**E**) Proteasomal activities in tissue lysates determined from Michaelis-Menten functions as in **Figure 3B** and color-coded for control (black), AD (yellow), PD (blue) donors; each measured in triplicate and from three donors per disease (see **Materials and Methods**). Proteasome concentrations were determined in **SFigure 11**. **(F)** A working model of TAAD formation triggered by aggregate presence. After their formation, TAADs engage and target aggregates, leading to their destruction. This enables proteasomes to be re-dispersed to resume their functions in regulating canonical proteostasis. Should aggregates become numerous or difficult to remove, this may instead lead to prolonged TAAD formation, preventing proteasomes re-dispersing to take part in cellular proteostasis and aggravating cell stress. This may eventually contribute to the progression of neurodegenerative disorders.

**Table 1.** List of antibodies used for immunofluorescence staining.

| Target Protein | Product code | Manufacturer | Species | Dilution |
| --- | --- | --- | --- | --- |
| P4D1 | 3936S | Cell Signaling | Mouse | 1:250 |
| FK2 | 04-263 | Merck Millipore | Mouse | 1:200 |
| K48-linked chains | ab140601 | Abcam | Rabbit | 1:250 |
| K63-linked chains | 05-1308 | Merck Millipore | Rabbit | 1:250 |
| UBQLN2 | 23449-1-AP | Proteintech | Rabbit | 1:250 |
| UBE2D | sc-166278 | Santa Cruz | Mouse | 1:250 |
| VCP | ab109240 | Abcam | Rabbit | 1:250 |
| HSP70/HSC70 | sc-32239 | Santa Cruz | Mouse | 1:250 |
| RAD23A | 24555S | Cell Signaling | Rabbit | 1:250 |
| RAD23B | 13525S | Cell Signaling | Rabbit | 1:250 |
| CHIP | ab109103 | Abcam | Rabbit | 1:250 |
| PSMB5 | sc-393931 | Santa Cruz | Mouse | 1:250 |
| PSMD14 | MA5-35818 | ThermoFisher | Rabbit | 1:200 |
| MAP2 | ab5392 | Abcam | Chicken | 1:1000 |
| CCT3 | 10571-1-AP | Proteintech | Rabbit | 1:200 |
| Goat anti-chicken | Alexa Fluor™ 405 | Abcam | ab175674 | 1:2000 |
| Goat anti-rabbit | Alexa Fluor™ 488 | ThermoFisher | A-11008 | 1:5000 |
| Goat anti-rabbit | Alexa Fluor™ 568 | ThermoFisher | A-11011 | 1:5000 |
| Goat anti-rabbit | Alexa Fluor™ 647 | ThermoFisher | A-21245 | 1:5000 |
| Goat anti-mouse | Alexa Fluor™ 488 | ThermoFisher | A-11001 | 1:5000 |
| Goat anti-mouse | Alexa Fluor™ 568 | ThermoFisher | A-11004 | 1:5000 |
| Goat anti-mouse | Alexa Fluor™ 647 | ThermoFisher | A-21235 | 1:5000 |

To address whether chaperones may assist in aggregate degradation, we stained and found HSC70/HSP70 with TAADs (**Fig.4E**), consistent with biochemical evidence of HSC70/HSP70-based aS disaggregases^31–33^ and that HSP70 likely interacts with proteasomes during oxidative stress for their reactivation^34^. Unexpectedly, TAADs did not appear to contain the hexameric AAA+ ATPase VCP/p97, which interacts with both proteasomes and HSP70 and implicated in targeting tau aggregates^35,36^ and Alzheimer’s disease^37^ (**SFig.6E**). Instead, TAADs co-localized with CCT3 (**SFig.6F**), a critical subunit of the TRiC/CCT, another AAA+ ATPase complex known to interact with proteasomes^38^. This suggests that TAADs contain multiple chaperone/refolding functions in addition to proteolytic activity.

UBQLN2 and RAD23A/B are shuttling factors^39^ that reversibly interact with proteasomes and are known to bind aggregates. We therefore examined whether TAAD formation would be mediated through these factors. UBQLN2 was reported to bridge between HSP70 and proteasomes to facilitate aggregate clearance^40^ and indeed stained positive in TAADs (**SFig.6G**). RAD23B and its close homolog RAD23A both co-localized with TAADs (**Fig.4F and SFig.6H**), potentially signifying redundancy between shuttling factors within TAADs. To validate our observations, we performed knock-down of all three shuttling factors and found that only RAD23B was critical for TAAD formation at 3 and 24 hrs (**Fig.4G-H** and **SFig.7**). RAD23B was also reported important in LLPS-foci formation^12^, suggesting one shared feature between the formation of TAAD and LLPS-foci. Possibly, UBQLN2 and RAD23A may have supporting roles whereas RAD23B is the essential shuttling factor in TAADs. These observations together confirm our postulation that TAADs would involve distinct components of the UPS and chaperone systems to specifically engage internalized aS aggregates^2^.

### Altered proteome and proteostasis impact electron transport and cell metabolism

We expected significant changes to proteostasis following treatment with aS aggregates and examined cell lysates at different timepoints by proteomics analysis (see also **Materials and Methods**). We identified a greater number of proteins in the pellet (**Fig.5A-D**; 3,540 ± 271, mean ± SD) than in the supernatant fractions (**SFig.8A-D**; 2,778 ± 285, mean ± SD). Sucrose treatment primarily affected the supernatant, with rapid remodeling of the cytosolic proteome suggested by enrichment of GO terms associated with translation and ribosome biogenesis (**SFig.8E**), while pellet fractions showed only limited perturbation (**Fig. 5A**). In comparison, aggregate treatment suggested increase in proteins in the pellet with altered abundance 3 hrs to 24 hrs post-treatment with aS, that had lasting effects at 48 hrs (**Fig 5B-D**). We observed differential abundance of several proteins encoded by genes linked to metabolism (e.g. PGK1, GMPS), oxidative stress (e.g. CAT), and mitochondrial function (e.g. COX7C). Functional gene enrichment analysis (GO) suggested a sustained stress response 24 and 48 hrs post aggregate treatment, with terms linked to altered metabolic and mitochondrial processes (**Fig.5E**). This observation may reflect the link between aS aggregates and mitochondrial damage that precede the cellular phase of PD^41^.

Some proteins found in the pellet may also have a propensity to condensate and thus potentially contribute to facilitating TAAD formation and interaction with aggregates. To examine this, we modified a protocol^42^ to enrich for aS condensates and stress granules and subjected these samples to proteomics analysis (**SFig.9A-C**). Several condensation-prone proteins were identified in this enriched fraction (e.g. G3BP1, KRAS), possibly suggesting involvement in TAAD formation (**SFig.9D**). Alternatively, TAAD formation may have been driven by the same mechanism that caused these proteins to also condensate. Intriguingly, GO analyses (**SFig.9E-G**) indicated elevated nuclear processes, revealing aggregate-induced stress at the genome level. Together, our results indicate significant and lasting effects following aggregate internalization reflected by altered protein abundance.

### Evidence for TAADs in pathology

We hypothesized that TAADs constitute an acute stress response to target aggregates and restore normal proteostasis, while prolonged TAAD formation may contribute to augmented disruption to physiological degradation processes leading to neurodegeneration and alpha-synucleinopathy. To demonstrate disease relevance, we repeated experiments designed in **Fig.1D** and showed that cells incubated with pathological aggregates from PD donors also induced TAAD formation at 3 hrs in the nucleus, followed by co-localization with aggregates in the cytosol at 24 hrs and finally disappearing at 48 hrs (**SFig.10A**). We also repeated the incubation with recombinant aS aggregates in cortical neurons differentiated from iPSCs that carry a pathological *SNCA* triplication mutation, which again saw TAAD formation and co-localization with incubated aS aggregates (**Fig.6A** and **SFig.10B**). These observations indicate that TAAD formation can be induced by proteotoxic stress of pathological origin.

Further validation was sought from immunofluorescence staining of tissue slices from a PD donor, which indeed identified strong proteasome staining that co-localized with aggregates in neurons (**Fig.6B**). Examination of the same number of fields-of-views in a neurologically healthy control tissue did not detect strong proteasome-nor aggregate staining (**Fig.6C**), suggesting that TAAD-like proteasome-rich bodies are found in pathology. Interestingly, we observed similar strong co-localizations between aggregates and proteasomes in neurons found in tissues from an Alzheimer’s disease (AD) donor (**Fig.6D**). It is tempting to consider that possibly the pathological implication of such TAAD-like formation potentially extends beyond PD. To support this idea, we examined whether proteasomal degradation activity would be maintained in pathological brain tissues, as found in **Fig.3B**. From the Michaelis-Menten functions, we found no difference in proteasomal catalytic efficiencies between control, PD, and AD donor tissues (**Fig.6E** and **SFig.11**). While we do not exclude the possibility that a minor fraction of proteasome particles that co-localized with aggregates may be compromised, our data demonstrate that bulk degradation activity in tissues remained comparable to that in healthy controls.

Overall, our observations together point towards a model where aS aggregates induce TAAD formation that results in sequestered proteasomes excluded from participating in regular proteostasis activities. Such acute formation of TAADs in healthy cells facilitates aggregate removal, followed by re-dispersion of proteasomes. In cells where the internalized aggregates are difficult to degrade or where aggregate load increases faster than their removal, TAAD formation may be prolonged and lead to compromised proteostasis (**Fig.6F**). This, we propose, may enable or contribute to further progression of the harmful effects associated with pathological aS aggregates.

## Discussion

TAADs are, to our knowledge, the first example of gel-like proteasome-rich condensates. Their appearance seems to primarily enhance degradation activity against aggregates, as suggested by the presence of chaperones, proteasome shuttling factors and ubiquitinating enzymes. We propose three purposes of TAADs. Firstly, TAAD enables increased interactions between proteasomes and other enzymes/co-proteins, thus facilitating concerted disaggregation/degradation of aggregates, as we postulated^2^. Secondly, their gel-like state may further prevent endogenous proteins from entering the condensate and thereby protects cytoplasmic proteins from being unintentionally degraded. Finally, by engaging with TAADs, aggregates and partially degraded fragments^22^ are shielded from engaging with other cellular mechanisms in a harmful way until they become fully degraded. It is therefore plausible that temporarily formed TAADs are favorably accelerating aggregate removal and indirectly alleviate proteotoxic stress responses in healthy cells. In neurodegenerative conditions as observed for alpha-synucleinopathies however, the role of TAADs, may be challenged by an increasing load of pathological aggregates and result in an eventually irreversible state.

## Acknowledgement

This work was supported by an Alzheimer’s Research UK Major Projects Grant [ARUK-PG2023A-025]; a Royal Society Research Grant [RG\R1\241116], and a Sir Henry Wellcome Research Fellowship [101585/Z/13/Z], all awarded to Y.Y.

## Author contributions

LS and MM designed and performed experiments and analyzed data, with support from MB and TC. LS prepared samples and MB analyzed mass spectrometry data with support from HW. YY directed the research and prepared the manuscript. All authors contributed to writing the manuscript.

## Materials and Methods

### Cell culture and imaging preparations

Generation of HEK293A cells expressing eGFP- or mEos-tagged PSMD14 and PSMB2 subunits from their respective genomic loci is described in detail in our companion paper^3^. Briefly, the coding sequence for eGFP or mEos3.2 was inserted by CRISPR-editing at the 3’-end between the last codon and stop codon. The eGFP or mEos sequence was flanked by codons for a linker (GlyGlySer×3) found between the proteasomal subunit and the fluorescent protein sequence, and sequence coding for a myc or flag affinity tag. This created four monoclonal cell lines expressing PSMD14-(GGS×3)-eGFP-myc, PSMB2-(GGS×3)-eGFP-flag, PSMD14-(GGS×3)-mEos-myc or PSMB2-(GGS×3)-mEos-myc, which were incorporated into respective proteasomal particles and of comparable activity as in unedited cells^3^.

Cells were maintained at 37 °C and 5% CO_2_ in DMEM (Sigma) supplemented with 10% (v/v) fetal bovine serum (Life Technologies). Glass coverslips (0.17 mm thickness, Thorlabs) were cleaned with Argon plasma for 1 hr before incubation with cells and mounted on the microscope. Prior to imaging, cells were washed three times and imaged in warm Fluorobrite (Thermofisher) pre-filtered through filters with 0.02 µm pore size. A metal chamber (Thermofisher) was used to secure the coverslip above the objective.

An iPSC line expressing triplication mutation of the *SNCA* gene (gift from Matthew Wood, Oxford) was differentiated into cortical-like neurons following a dual-SMAD inhibition protocol^43^, plated onto imaging coverslips and matured up to >100 days-in-vitro prior to fixation and immunofluorescence staining.

### Immunofluorescence staining

Cells were grown on glass coverslips (25 mm diameter, VWR) in a 6-well plate until the desired confluency was reached or the appropriate treatment was carried out. We attempted several published protocols for cell fixation and optimized the following protocol to retain the highest level of TAADs at 24 hrs. Paraformaldehyde (8%) was used in a 1:1 dilution with growth media for direct cell fixation on the coverslip, which was incubated on the bench for 15 min. This procedure partially retained TAADs at 24 hrs, with a ∼7-fold reduction in their level and ∼2-fold size enlargement by area (**SFig.5**). The same fixation protocol also reduced the number of LLPS-foci per cell by ∼140-fold and increased their area by ∼2-fold. Attempts to retain TAADs at 3 hrs using this protocol or using different concentrations of PFA, by addition of 0.2% glutaraldehyde to the fixative solution, or fixation with methanol were all unsuccessful (data not shown).

Following fixation, cells were washed with PBS for 3× 5 min and permeabilized using ice-cold methanol at −20 °C for 5 min. Cells were then washed with PBS 3× for 5 min prior to blocking in 10% goat serum diluted in PBS for 1 hr. Primary antibodies were diluted in 5% goat serum diluted in PBS and incubated with cells overnight at 4 °C. Cells were then washed with PBS 3× for 5 min followed by secondary antibody incubation in 5% goat serum for 1 hr. Prior to imaging, a final 3× 5 min PBS wash step for was carried out before mounting on the microscope. Primary and secondary antibody information are shown in **Table 1**.

### Inhibitors and reagents

Cells were treated with the following inhibitors and dilutions: colcemid (Merck), MG262 (Stratech), or TAK-243 (Selleck Chem) at concentrations stated in the figures, all prepared in DMSO. After the stated incubation period, cells were taken out from the incubator and washed three times using PBS before imaging. Stock 1,6-hexanediol (Merck) was added to the media reaching 5% final concentration and incubated with cells with pre-formed foci for up to 1 hr. For Me4BodipyFL-Ahx3Leu3VS (Bio-Techne Ltd) treatment cells were incubated at 1 µM final concentration for 1 hr at 37 °C before wash step and imaging.

### siRNA knockdown

PSMD14-eGFP cells were grown to 90% confluency in 6-well TC-treated dishes (Corning). siRNAs targeting human RAD23A, RAD23B and UBQLN2 were purchased from Merck (EHU027621, EHU145881 and EHU029491, respectively). These siRNAs were provided as heterogeneous mixtures designed to target the same mRNA sequence and were re-suspended in TE buffer (10 mM Tris-HCl pH 8, 1 mM EDTA). Lipofectamine™ 2000 (ThermoFisher) was used to transfect each well with 4 µg siRNA. After 24 hrs, cells were re-plated onto coverslips and imaged 72 hrs post-transfection. Knockdown efficiency was verified by immunoblotting cell lysates. siRNA targeting Kif11 (EHU019931) was used as the positive control for transfection efficiency and monitored for rounding of cells in response to mitotic arrest. As a non-targeting siRNA control, cells were transfected with siRNA for Renilla Luciferase (RLUC) (Merck).

### TIRF and HILO instrument and imaging 2D

Imaging was performed using a home-built system^1^ using the ECLIPSE Ti2-E inverted microscope (Nikon) with CFI Apochromat TIRF 100XC NA 1.49 Oil objective (Nikon) and 1.5× tube lens. The change from HILO to TIRF mode was achieved by altering the beam angle using an E-TIRF arm (Nikon). A short pulse of 405 nm was used to photoconvert mEos, which was subsequently excited using the 561 nm laser. Images were recorded on a Prime95B sCMOS camera, giving pixel size at 73 nm after 150× magnification.

### SMLM reconstruction

SMLM images in 2D were reconstructed using Matlab scripts described^44^. Images of TetraSpeck™ Microspheres (Life Technologies) deposited on coverslips were collected in all four laser channels alongside each multi-color experiment to allow for correction of chromatic aberrations between channels for alignment. SMLM in 2D achieved 21 ± 1 nm planar resolution^1^.

SMLM images in 3D were reconstructed using the ThunderSTORM plug-in for ImageJ^45^. Calibration files of the TetraSpeck™ Microspheres were produced by recording stacks of images along the z-axis with the asymmetric lens installed in the Optosplit II. The lens distorted the point spread function (PSF) recorded from each fluorescent particle relative to position of the particles above or below the focal plane. The resultant reference curves were then used to calculate the axial positions of fluorophores inside cells or on the coverslip surface. This 3D SMLM approach achieved FWHM of 33 nm, 42 nm, and 84 nm for x, y, and z directions, respectively^1^.

### Quantitative analysis of imaging data

Images were analyzed using custom-written Matlab codes to quantify the number and size of TAADs, LLPS-foci and aggregates in cells. For each field of view, we typically collect and average 100 frames at 20 Hz. The average fluorescence signals are then background-subtracted using a rolling ball and a Gaussian filter, through which the threshold for each particle is determined. Cell boundaries were identified using a rolling ball filter for GFP and a threshold applied to identify TAADs. LLPS-foci were analyzed by a similar script that also accounted for the proximity and overlap between each condensate type by preferentially identifying local intensity maxima within the cell boundaries over detecting foci using a global intensity threshold. Foci number, size, and eccentricity were then quantified using the masks detected by the Matlab scripts.

Co-localization analysis was performed using the ComDet plugin (v.0.5.5) for ImageJ software. Image from multiple channels were imported into ImageJ and channels merged. Co-localization is defined as when the centers of each spot in two channels (for GFP and AF647) are within two pixels. The approximate particle size for TAADs and aggregates was set to 10 pixels and 5 pixels, respectively and intensity thresholds were manually determined for each set of images, typically in the range of 15-20. Random co-localization was determined by rotating images in the AF647 channel 180° and repeating the analysis with the same parameters.

All our codes are available for download on https://github.com/ye-team-organisation/TAAD-analysis.

### FRAP analysis

FRAP experiments were performed using a Leica TCS SP8 laser-scanning microscope equipped with HC PL APO oil objective giving 63× magnification and NA 1.40 (Leica Microsystems, 11506350). Circular ∼1 μm^2^ regions of interest (ROI) were identified in cells and 10 images (1 frame/s) taken prior to bleaching. ROIs were then bleached using a 1 s pulse of the 488 nm laser at full power (65 mW) and recovery was monitored every 1 s for 500 s. Data was corrected for photobleaching occurring during imaging and normalized to pre-bleach fluorescence. Plotting and curve fitting was carried out using GraphPad Prism 10 software.

### Preparation of recombinant aS

Recombinant aS was purified, labeled and aggregated as described previously^1^. Plasmids encoding full-length wild-type or Q24C aS in pT7-7 vector were transformed into BL21 (DE3) pLysS cells and expressed with 1 mM IPTG induction. Cell lysate was cleared by sequential biochemical and high-speed centrifugation steps and loaded onto a HiTrap CaptoQ column for separation. Fractions containing the protein were pooled and dialyzed overnight against 25 mM Tris (pH 8.0 at 4 °C) buffer. Labelling with AlexaFluor 647 maleimide was done in >2:1 excess of the fluorophore at room temperature for 1 hr followed by size exclusion chromatography to remove unlabeled fluorophore. Prior to aggregation, we mixed wild-type aS with AF647-labeled aS monomers in 19:1 ratio to allow formation of fibrils incorporating the labeled monomers. The aggregation reaction lasted for 72 hrs under 200 rpm agitation at 37 °C.

### Protein gel electrophoresis and immunoblotting

Samples were quenched with LDS loading buffer (ThermoFisher) for separation by SDS-PAGE on 4-16% Bolt gels (ThermoFisher). We used semi-dry transfer of separated proteins onto a PVDF membrane followed by standard immunoblotting procedures^1^ using antibody combinations and dilutions as specified in **Table 2**.

**Table 2.** List of antibodies used for immunoblotting.

| Target | Product code | Manufacturer | Species | Dilution |
| --- | --- | --- | --- | --- |
| PSMB2 | sc-58410 | Santa Cruz | Mouse | 1:1,000 |
| PSMD14 | MA5-35818 | ThermoFisher | Rabbit | 1:1,000 |
| PSMB5 | sc-393931 | Santa Cruz | Mouse | 1:1,000 |
| MJFR1 | ab138501 | Abcam | Rabbit | 1:1000 |
| UBQLN2 | 23449-1-AP | Proteintech | Rabbit | 1:250 |
| RAD23A | 24555S | Cell Signaling | Rabbit | 1:250 |
| RAD23B | 13525S | Cell Signaling | Rabbit | 1:250 |
| Alexa Fluor™ 488 | A-11008 | ThermoFisher | Goat anti-rabbit | 1:10,000 |
| Alexa Fluor™ 647 | A-21235 | ThermoFisher | Goat anti-mouse | 1:10,000 |

To determine proteasome concentration in lysates, we created a standard curve from purified 26S proteasome holoenzymes of known concentration, from which the concentration of PSMB5 in cell lysates was determined from immunoblotting. Cell or tissue lysates were loaded and immunoblotted on the same PVDF membrane with the same antibody and the proteasome concentrations in respective lysate was determined by densitometry analysis against a standard curve of band intensity versus the known concentrations.

### Measurement and analysis of enzymatic reactions

Succinyl-Leu-Leu-Val-Tyr-7-amino-4-methylcoumarin (Suc-LLVY-AMC, Abcam) in DMSO was serially diluted to the stated concentrations and incubated with cell or tissue lysates, which was keep at constant final concentrations. Initial rates of reaction were determined by first measuring gain in fluorescence over time until a plateau maximum is reached, which follows a negative single-exponential function, and then taking the first order differential value at origin. We have previously demonstrated calculation of proteolytic enzyme kinetics analysis using this approach^46^, which does not rely on determining the absolute fluorescence values across a large concentration range. The initial rates were then plotted against substrate concentration and fitted with the Michaelis-Menten function:

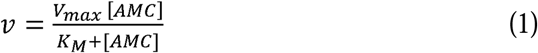

where *v* is the rate of reaction, *V_max_* is the max reaction rate, *K_M_* is the Michalis constant and [AMC] the concentration of substrate. The turnover number, *k_cat_*, is calculated from

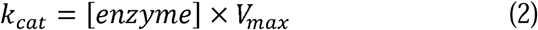

where [enzyme] is determined by densitometry analysis of immunoblotting against proteasome subunits.

To compare the catalytic efficiency between different enzymes or enzymes under different conditions, we calculated specificity constant as follows

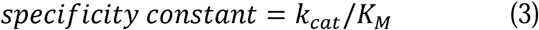

expressed in M^-1^s^-1^.

### Enzyme kinetics measurements of proteasomes in lysates

Untreated PSMD14-eGFP cells, cells incubated with sucrose, with aggregates or flash-frozen tissues (**Table 3**) from Multiple Sclerosis and Parkinson’s Tissue Bank were resuspended in lysis buffer (50 mM Tris pH 7.4 at 4 °C, 5 mM MgCl_2_, 2 mM ATP) and lysed by a motorized pestle mixer. Supernatants were cleared by a benchtop centrifuge at 15,000 rpm for 20 min at 4 °C. Concentrations were measured by Bradford assay and 20 µg of lysates was taken out to dilute in assay buffer (50 mM Tris pH7.4, 150 mM NaCl, 5 mM ATP) to 100 µl final volumes for clear-bottomed 96 well-plate (Greiner). Suc-LLVY-AMC (Bio-Techne) stocks were diluted directly in wells to the indicated concentration (5-400 µM). Fluorescence was measured very 90 s for 25 hrs at 37 °C using a CLARIOstar microplate reader (BMG Labtech) with λ_Em_ = 350 ± 15 nm and λ_Ex_ = 440 ± 20 nm. To calculate *k_cat_* values, final lysate proteasome concentrations were calculated by immunoblotting and densitometry analysis against purified proteasomes of known concentrations.

**Table 3.**
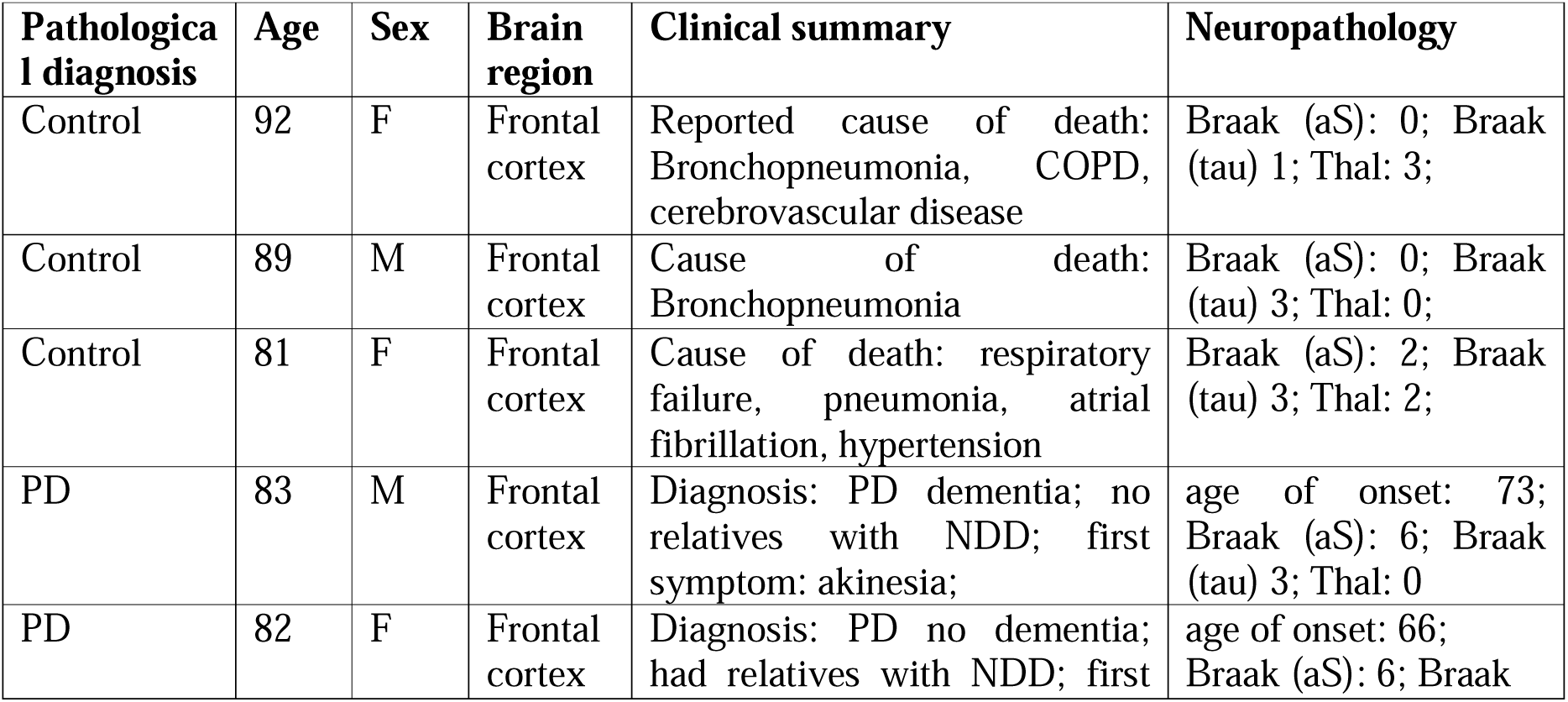
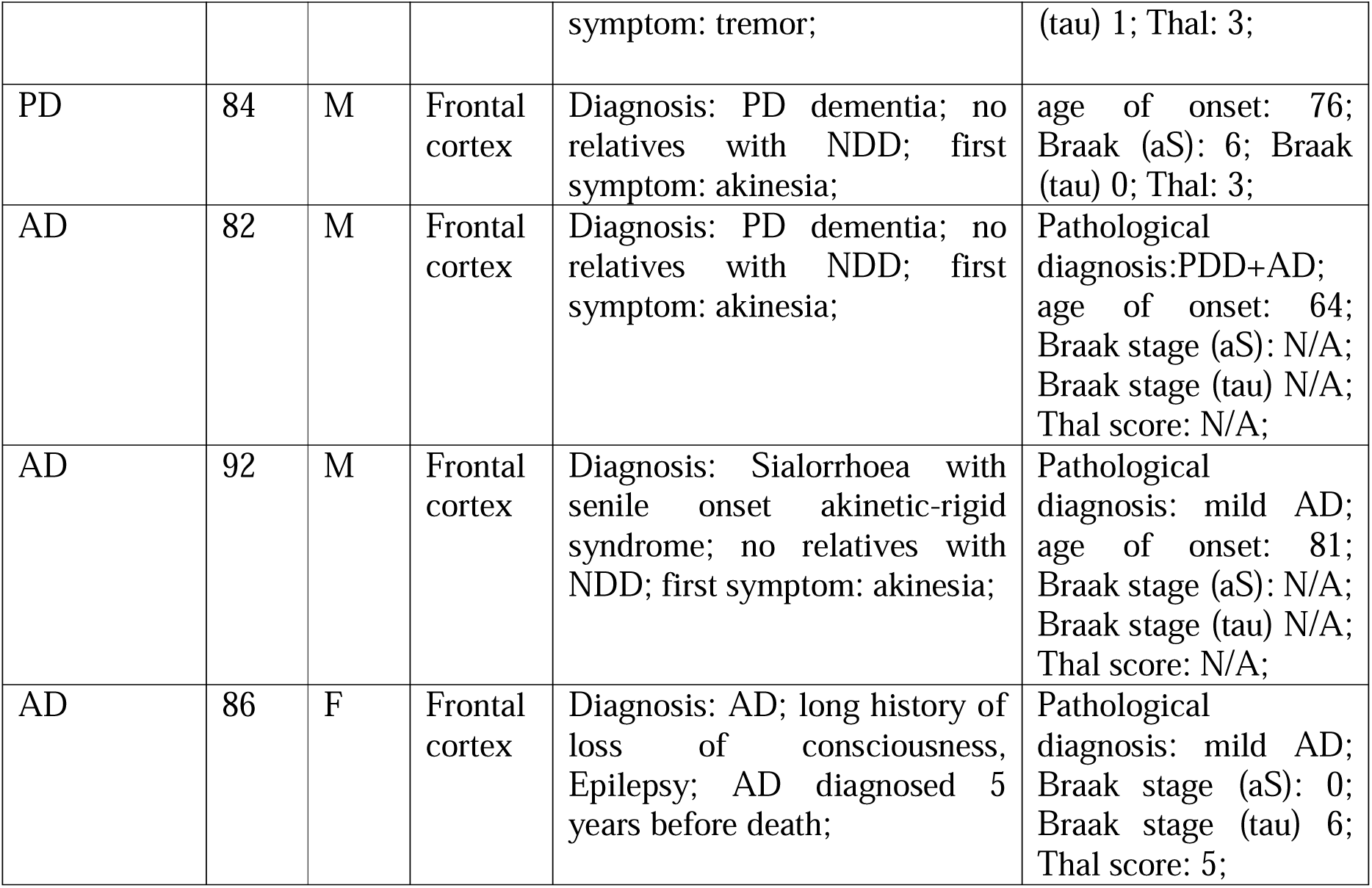
Human brain tissues used for proteasome Michaelis-Menten kinetics measurements.

### Sample preparation and mass spectrometry

Cells grown in 6-well plates (Corning) and either left untreated, incubated with sucrose or aS aggregates. At the defined timepoints, cells were washed 3× in PBS and collected by scraping in lysis buffer for whole cell lysate proteomics (50 mM Tris pH 7.4 at 4 °C, 5 mM MgCl_2_, 2 mM ATP) or for TAAD enrichment (lysis buffer + 0.5% NP-40). For lysate proteomics, cells were lysed by homogenization using a motorized pestle mixer (VWR) for 30 s. Lysates were then centrifuged at 15,000 rpm for 20 min at 4 °C to separate soluble and insoluble fractions. Soluble fractions were transferred to new tubes and insoluble fractions resuspended in 20 µl lysis buffer. Concentrations of fractions were determined by Bradford assay, and samples were flash frozen and stored at −80 °C. Cell treatment and sample collection was repeated 3× with a maximum of two passages between replicates. Once all replicates had been collected samples were thawed and 20 µg of each sample aliquoted to new tubes for subsequent processing.

A modified protocol (**SFigure 9A**) for isolating stress granule cores was used for enrichment experiment^42^, where cells were lysed by repeatedly passing through a 25G needle followed by centrifugation at 1,000× g for 5 min at 4 °C. The cleared supernatant was subjected to 18,000× g centrifugation for 20 min and the pellet resuspended to repeat this procedure twice. The final resuspended pellet was centrifuged at 850× g for 2 min, and this pellet was used for mass spectrometry analysis.

For mass spectrometry preparation, each sample was reduced, alkylated and digested at 37 °C in a single-pot reaction using an automated workflow on an Andrews^+^ Pipetting Robot (Waters) in which the sample is combined with 40 mM chloracetamide, 10 mM TCEP and 0.1 μg sequencing grade modified porcine trypsin (Promega) in 60 μl of 100 mM ammonium bicarbonate and held at 37 °C for 16 hours. Subsequently, samples are cooled to 4 °C until before being acidified with 0.5% trifluoroacetic acid (TFA). Sample were desalted using HLB 30 μM μElution Plates (Waters) using a vacuum manifold. The HLB plate was first primed by 3× 100 μL of 80% acetonitrile and the matrix equilibrated with 3× 100 μL of 0.1% TFA. Each sample was then passed through and washed with 3× 100 μL 0.1% TFA before being eluted with 3× 50 μl of 60% acetonitrile into a Waters QuanRecovery plate. Samples were dried completely in a Speedyvac (Eppendorf) at 60 °C (∼2 hours) and stored at −80 °C. All steps were performed using an Andrew+ Pipetting Robot (Waters).

For acquisition, each sample was thawed on bench and dissolved in 200 μl of 0.1% formic acid (FA) for 10 minutes sonicating followed by centrifuging for 1 minute at 1000 ×g. Samples were then loaded into a UPLC system (M-Class Acquity, Waters) and 4 μl was separated across a Kinetex™ 150 × 0.3mm 2.6 μm XB-C18 column (Phenomenex) interfaced with a high-performance Q-TOF mass spectrometer or MS (ZenoTOF 7600, Sciex). Chromatography was performed at 10 μl/minute with 3% Buffer A (water with 0.1% FA) and 3% Buffer B (acetonitrile with 0.1% FA) increasing to 30% Buffer B between 1 and 21 minutes. The column was then washed with 80% organic solution for 3 minutes before equilibrating for 5 min prior to the next injection. The MS was run in positive mode with a zenoSWATH method comprising 85 variable windows with ionization scheduling between 2 and 27 minutes. MS data was acquired over 400-1500 mz with an accumulation time of 0.1 s. MS/MS was acquired over 140-1800 mz with an accumulation time of 0.013 s and dynamic collision energy enabled.

### Proteomics data analysis

MS files (.wiff) were imported into DIA-NN version 2.2 and searched using an *in-silico* predicted library (created in DIA-NN) from the Swissprot Reference Human Proteome (downloaded 8^th^ September 2025) supplemented with PSMD14-eGFP (570 Aa, 63359.33Da; see **Supplementary Sequence File**). Output was set to be filtered to 1% FDR (default) with the maximum number of variable modifications set to 2, methionine variably oxidized and all cysteines assumed carbamidomethylated. Match between runs was enabled.

Differential protein abundance was analyzed in R (R version 4.5.1), using DEqMS^47^. Protein group intensities were loglJ-transformed, filtered for missingness by protein, and median-centered across samples. For whole-cell (supernatant, pellet) and enriched experiments, proteins were required to have ≥ 2 quantified values in each condition for a given comparison (i.e. treatment vs control). Statistical significance was assessed via moderated t-test with empirical Bayes variance shrinkage (limma) and adjusted for variance on the peptide level implemented through the DEqMS package^48^. P-values were corrected for multiple testing using the Benjamini–Hochberg method to control the false discovery rate (FDR). Differentially abundant proteins were defined at |loglJFC| > loglJ(1.5) and DEqMS (spectrally adjusted) p-value < 0.05, with and without FDR correction for enriched and whole cell (supernatant, pellet) datasets, respectively.

Functional enrichment was performed on significant proteins using clusterProfiler::enrichGO (ontology: BP; OrgDb = org.Hs.eg.db) and ReactomePA::enrichPathway, using custom backgrounds defined as all filtered proteins in that comparison and a minimum overlap of 2 genes with the background. For enriched data, proteins present exclusively in the treated/control (defined as present in ≥ 4/5 replicates and 0/5) were combined with differentially abundant proteins for to generate the enrichment input. The top 15 terms, ranked by FDR, were reported. For whole-cell data, the top 10 GO terms per treatment with FDR < 0.05 were selected. To minimize redundancy in this selection, we first pruned overlapping gene sets (Jaccard index ≥ 0.8), retaining the lowest FDR term, then clustered semantically similar terms with GOSemSim^49^ and rrvgo::reduceSimMatrix (threshold ≥ 0.7), and lightly shortened representative labels.

Full results are provided in **Supplementary Tables 1-3** and all analysis scripts are available at https://github.com/ye-team-organisation/TAAD-analysis.

The mass spectrometry proteomics data have been deposited to the ProteomeXchange Consortium via the PRIDE^50^ partner repository with the dataset identifier PXD068227.

### Human Tissues

Flash-frozen and paraffin-embedded 7 µm tissue sections from the frontal cortex of control, PD and AD donors (**Table 4**) were obtained from the Multiple Sclerosis and Parkinson’s Tissue Bank or MRC Neurodegenerative Diseases Brain Bank. Following standard protocols, tissue sections were rehydrated and dewaxed. Prior to staining, pre-treatment included blocking of endogenous peroxidases with 3% hydrogen peroxide and 80% formic acid. Sections were then incubated overnight with 1:250 dilution of antibodies against PSMB5, MAP2 (see **Table 1**) and 1:1000 dilution of Amytracker-630^1^ (AT630). Primary antibodies were visualized using secondary antibodies in 1:1000 dilution. Slides were dehydrated and cleared prior to mounting.

**Table 4.** Human brain tissues used for immunostaining.

| Pathological diagnosis | Age | Sex | Post-mortem interval (hrs) | Brain region | Neuropathology |
| --- | --- | --- | --- | --- | --- |
| Control | 73 | F | 27 | Temporal cortex | Very mild neuronal loss in the cortex. Sparse predominantly diffuse type of B-amyloid positive plaques without labelling of the blood vessels by B-amyloid. Only occasional tau positive pretangles are seen.<br>The brain revealed only very early ageing changes in forms of tauopathy affecting the amygdala, entorhinal cortex, and subiculum. The tangle stage is consistent with stage I according to BNE classification. There was no evidence of Alzheimer's disease, a-synucleinopathy, or TDP-43 proteinopathy. |
| PD | 76 | M | 66 | Temporal cortex | Neuropathological report missing from MRC Neurodegenerative Diseases Brain Bank |
| AD | 82 | M | 12 | Frontal cortex | Alzheimer-type tau pathology, Braak Stage I |

## Supplementary Figures

**SFigure 1.** Representative images of HEK293A cells expressing PSMB2-eGFP from their genomic loci. **(A)** PSMB2-eGFP cells left untreated, (**B**) incubated with sucrose or **(C)** with CaCl_2_ and imaged by HILO microscopy as in **Fig.1A-C**. **(D)** PSMB2-eGFP cells incubated with 1 µM AF-647-labeled aS aggregates (red) for 3, 24 or 48 hrs as in **Fig.1D**. (**E**) Quantification of TAADs in untreated PSMB2-eGFP cells following 3 hrs (0.10 ± 0.36, n=71 cells), 24 hrs (0.27 ± 0.69, n=70 cells), and 48 hrs (0.14 ± 0.52, n=61 cells). Data are mean ± SD and were analyzed by t-test. (**F**) Representative images of PSMD14-EGFP cells ectopically expressing mCherry-LaminB1 (Addgene 55069), which enabled definition of nuclear envelope in cells under different conditions and permitted calculation of nuclear versus cytoplasmic fractions in **Fig.1E**. **(G)** Quantification of the percentage of cells containing aggregates at the different timepoints in **D** (3hrs: 81.5 ± 2.6, n=262 cells; 24hrs: 88.2 ± 2.7, n=209 cells; 48hrs: 37.2 ± 1.5, n=229 cells). **(H)** Quantification of the percentage of PSMD14-eGFP cells containing proteasome condensates (untreated: 3.7 ± 3.3, n=305 cells; sucrose= 100.0 ± 0.0, n=84 cells; 3hrs: 72.9 ± 6.8, n=262 cells; 24hrs: 74.9 ± 8.9, n=209 cells; 48hrs: 23.6 ± 8.5, n=229 cells) Error bars represent mean ± SD of three independent experiments. All concentrations represent final concentrations in solution. Scale bars represent 5 µm in all images.

**SFigure 2.** TAADs are not formed following incubation with monomeric aS or the aggregation buffer. (**A**) Representative images of PSMD14-eGFP cells treated with aS monomer at different timepoints and (**B**) quantification of the number of foci following incubation with aS monomer at 1 μM for 3 hrs (0.23 ± 0.60, n=79 cells), 24 hrs (0.11 ± 0.31, n=76 cells) or 48 hrs (0.18 ± 0.42, n=68 cells) compared with cells without incubation (0.053 ± 0.29, n=57 cells). (**C**) Representative images of PSMD14-eGFP cells incubated with aggregation buffer and **(D)** quantification of foci detected at 3 (0.09804 ± 0.3608, n=51 cells), 24 (0.2667 ± 0.6856, n=60 cells) and 48 hrs (0.1404 ± 0.5154, n=57 cells) compared with untreated control (0.03922 ± 0.1960, n=51 cells). Data are mean ± SD and analyzed by t-test.

**SFigure 3.** TAADs are distinct to LLPS-foci. **(A)** *Left*: representative images of PSMD14– eGFP cells containing LLPS-foci following incubation with sucrose and subsequently treated with 5% 1,6-HD for 0 and 60 min. *Right*: quantification of foci number per cell at 0 min (mean=2743 ± 1194, n=71 cells), 15 min (mean=54.3 ± 43.8, n=97 cells), 30 min (mean=65.7 ± 50.7, n=100 cells), 45 min (mean=63.5 ± 47.0, n=91 cells) and 60 min (mean=65.8 ± 50.2, n=93 cells) post treatment. (**B-C**) Quantification of FRAP experiments fitted to single-exponential functions for **(B**) untreated nucleoplasm and (**C**) sucrose-treated cells, performed as in **Figure 3E-F**. Results are mean ± SD from three independent experiments.

**SFigure 4.** Measurement of cellular proteolytic activity of proteasomes. **(A)** Representative images of resting PSMD14-eGFP cells (control) stained with Me4BodipyFL-Ahx3Leu3VS activity reporter as in **Figure 3A**. **(B)** Immunoblotting against PSMB5 in the lysates used for Michaelis-Menten kinetics in **Figure 3B**. All lysates were first measured by Bradford assay for loading adjustment and finally determined against a standard curve from band intensities of purified 26S proteasome holoenzyme^22^ at increasing concentrations. **(C)** The number ± SD of RP or **(D)** CP found in LLPS-foci (mean= 40 ± 10 and 12 ± 4, respectively) and TAADs (at 3 hrs: 70 ± 20 and 40 ± 10, respectively; and at 24 hrs: 300 ± 100 and 80 ± 30, respectively) were determined from quantitative SMLM imaging in CRISPR-engineered cells expressing PSMD14-mEos or PSMB2-mEos, respectively. For each condition, 50 TAADs were selected from cells used for **Figure 1** and **SFigure 1**. **(E)** The number of proteasome particles calculated present in TAADs or LLPS-foci was determined from the relative fluorescence intensity of single mEos fluorophores in PSMB2-or PSMD14-mEos cell lines. Total fluorescence emission measured from each foci was then calculated individually using the average mEos step size. See **Materials and Methods** for details. Scale bars represent 5 μm and 1 μm on the left and right, respectively.

**SFigure 5.** Fixation alters the appearance and abundance of TAADs and LLPS-foci. (**A**) Representative image of PSMD14-eGFP cells incubated with 200 mM sucrose for 45 min and imaged live or following fixation by protocol optimized as detailed in **Materials and Methods**. (**B**) Quantification of LLPS-foci per cell following sucrose incubation and imaged in live cells (2846 ± 1214, n=42 cells) or after fixation (18.3 ± 21.7, n=47 cells). **(C)** Quantification of LLPS-foci areas in live (0.16 ± 0.03, n=1272) or fixated cells (0.37 ± 0.10, n=731) as in **B**. **(D)** PSMD14-eGFP cells incubated with aS aggregates for 24 hrs imaged live or fixed and presented as in **A**. Quantification of (**E**) TAADs per cell before (7.20 ± 7.80, n=49 cells) or after fixation (0.76 ± 1.77, n=50 cells) incubated with aS aggregates for 24 hrs. (**F**) Quantification of area covered by TAADs in live (0.44 ± 0.15, TAADs=50, n=50 cells) or fixed cells (0.86 ± 0.44, TAAD=63, n=49 cells). (**G**) Quantification of the aggregate number per cell before (28.4 ± 19.0, n=50 cells) or after fixation (24.8 ± 14.7, n=49 cells). Data are mean ± SD and were analyzed by two-tailed Mann-Whitney U-test. Scale bars represent 5 μm.

**SFigure 6.** Examining protein components in TAADs. PSMD14-eGFP (green) cells were incubated with aS aggregates (red) for 24 hrs and fixed as described in **Materials and Methods**. Representative images selected from immunostaining performed with respective antibodies (blue) against **(A)** ubiquitin (clone P4D1), **(B)** Lys63-linked ubiquitin chains, **(C)** UBE2D, **(D)** CHIP, **(E)** VCP/p97, (**F**) CCT3, (**G**) UBQLN2, (**H**) RAD23A. Line profiling of respective cell sections as indicated by white dashed lines are color-coded accordingly. Representative images from at least three biological repeats, presented as in **Figure 4A**. Scale bars = 5 µm.

**SFigure 7.** Knockdown of UBQLN2 and RAD23A/B expression. **(A)** Kif11 knockdown triggers mitotic arrest and served as a positive control for siRNA transfection. Phenotypic examination showed rounded cells 48 hrs post-transfection, confirming mitotic arrest. Scale bars represent 1 mm. **(B)** Representative images of resting PSMD14-eGFP cells (no siRNA) or 48 hrs post transfection transfected with siRNA against RLUC, UBQLN2, RAD23A or RAD23B. **(C)** Number of foci per PSMD14-eGFP cell at resting state (0.10 ± 0.37, n=96 cells) or after transfection of siRNA against RLUC (0.33 ± 0.60, n=126 cells), UBQLN2 (0.21 ± 0.54, n=113 cells), RAD23A (0.39 ± 0.77, n=125 cells) or RAD23B (0.21 ± 0.46, n=104 cells). Data are mean ± SD and were analyzed by t-test. **(D-F)** Immunoblotting against **(D)** UBQLN2, **(E)** RAD23B or **(F)** RAD23A in untransfected cell lysates or lysates of 72 hrs post transfection of respective siRNA to verify knockdown at the protein level. Ponceau S staining was performed as a loading control.

**SFigure 8.** Analysis of proteomic changes in the supernatant after treatment with **(A)** sucrose, or **(B)** 3 hrs, **(C)** 24 hrs and **(D)** 48 hrs after aS aggregate incubation in PSMD14-eGFP cells (n=3 biological replicates). Results are presented as in **Figure 5** with the top 15 candidates labeled by gene name. See also **Supplementary Table 2**. **(E)** Heatmap of gene ontology (GO Biological Processes) over-representation of up-regulated proteins across all treatments. Displayed as top 10 GO terms with FDR < 0.05 from each treatment, curated to minimize term redundancy. Colored according to −log_10_(FDR) (Benjamini–Hochberg adjusted p-values).

**SFigure 9.** Analysis of proteomic changes in the enriched fraction following 24 hrs incubation with aS aggregates in PSMD14-eGFP cells (n=5 independent repeats). **(A)** Workflow of stepwise centrifugation leading to enrichment of stress granule cores in MS2, modified from a previous protocol^42^. **(B)** Silver-staining of untreated or cells incubated with aS aggregates for 24 hrs prior to mass spectrometry analysis. **(C)** Immunoblotting of samples at each step of enrichment against 20S proteasomal subunit PSMB5 (*left*) or aS (*right*), showing gradual concentration of the pellet fractions. (**D**) Differentially expressed proteins identified (Control: 1325 ± 105 proteins; 24 hrs: 1492 ± 91 proteins) are presented as in **Figure 5**. All hits with FDR-corrected, spectrally adjusted p-values < 0.05 and |logFC| > 1.5 are labeled by gene name. Functional enrichment analysis of the top 15 Reactome pathways **(E)** and GO Biological Process terms for upregulated **(F)** and downregulated **(G)** genes. Displayed by gene ratio, with dot size indicating hit count and colored according to the FDR-adjusted p-value.

**SFigure 10.** Validation of TAADs in iPSC-differentiated cortical neurons and with protein aggregates of pathological origin. **(A)** Representative images of PSMD14-eGFP cells treated with PD donor-derived soluble aggregates at 1 μM for 3, 24 or 48 hrs. Aggregates were stained with Amytracker-630^1^. Scale bars represent 5 μm. **(B)** Representative images of an iPSC-differentiated cortical neuron examined by immunostaining using anti-PMSD14 (blue), anti-MAP2 (white), anti-PSMB5 (green) antibodies and AF647-labeled aS aggregates (red), each shown in a single-color channel. See **Figure 6A** for co-localization between PSMB5 and aS aggregates.

**SFigure 11.** Proteasome concentrations in post-mortem tissues. **(A)** Tissue lysates from three each of control and Parkinson’s disease (PD) and **(B)** three each of dementia with Lewy bodies (not used for this work) and Alzheimer’s disease (AD) donors are immunoblotted with anti-PSMB5 and used for calculations in **Figure 6E**. Proteasome concentrations are then determined against the purified 26S proteasome holoenzyme loaded and calculated as in **SFigure 4B**.

## Supplementary Videos

Supplementary Videos can be found on the online version of this manuscript and on youtube.com (https://www.youtube.com/playlist?list=PLgc4yUNxF7D371orab4jtLxDKxPvUkEzR)

**SVideo 1**. Reconstruction of LLPS-foci as described in **Figure 2G**.

**SVideo 2**. Reconstruction of TAAD at 3 hrs as described in **Figure 2G**.

**SVideo 3**. Reconstruction of TAAD at 24 hrs as described in **Figure 2G**.

**SVideo 4**. Reconstruction of TAAD at 24 hrs with aS aggregate as described in **Figure 2H**.

**Supplementary Table 1**. Protein list from pellet of cell lysate.

**Supplementary Table 2**. Protein list from supernatant of cell lysate.

**Supplementary Table 3**. Protein list from enrichment procedure.

**Supplementary Sequence File**. Sequence of PSMD14-(GGS×3)-eGFP-myc in fasta format.

