## Supplementary Figures S1-S11 for "A novel type of gel-like proteasome condensate induced by toxic protein aggregates"

### Supplementary Figure 1

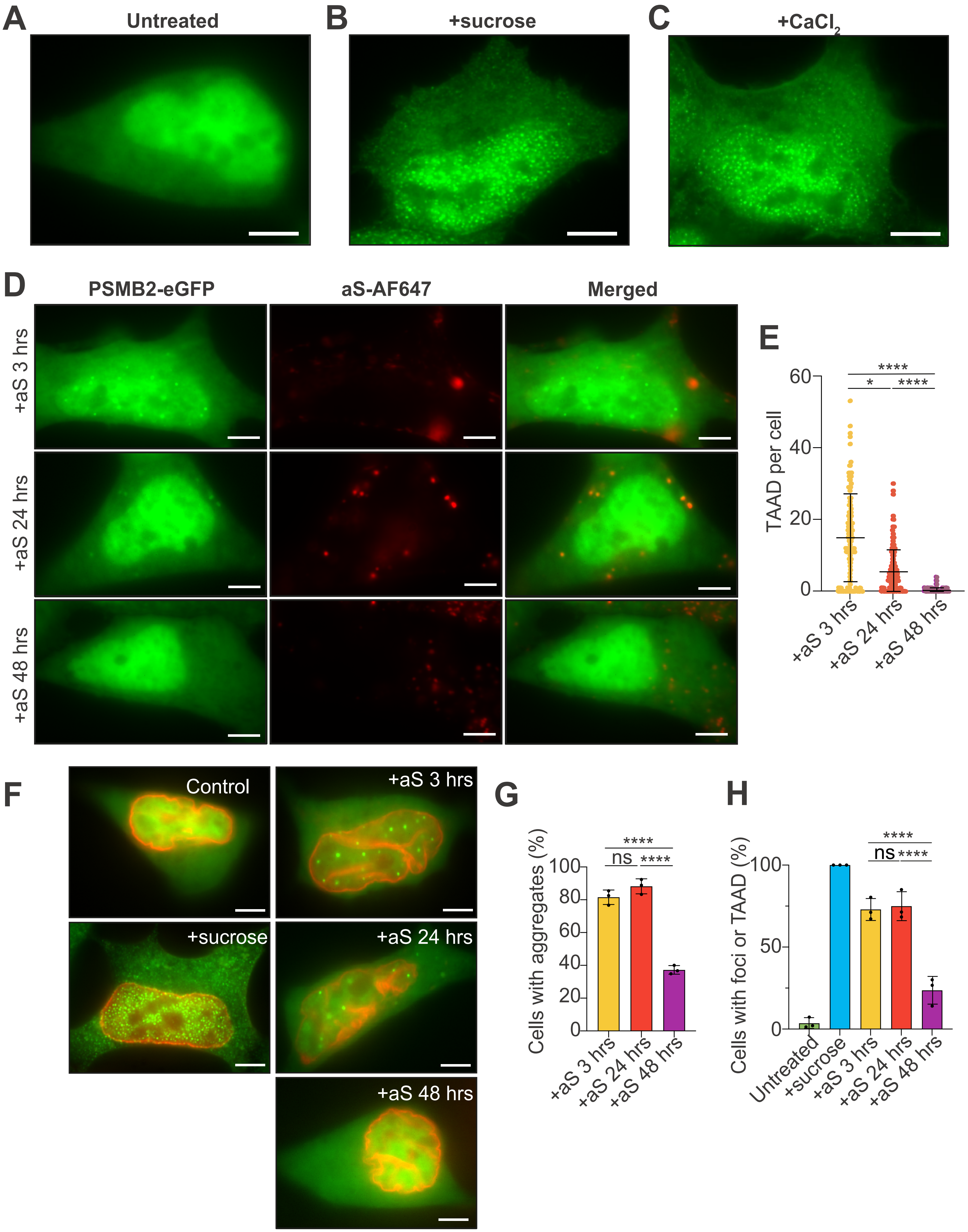

### Supplementary Figure 2

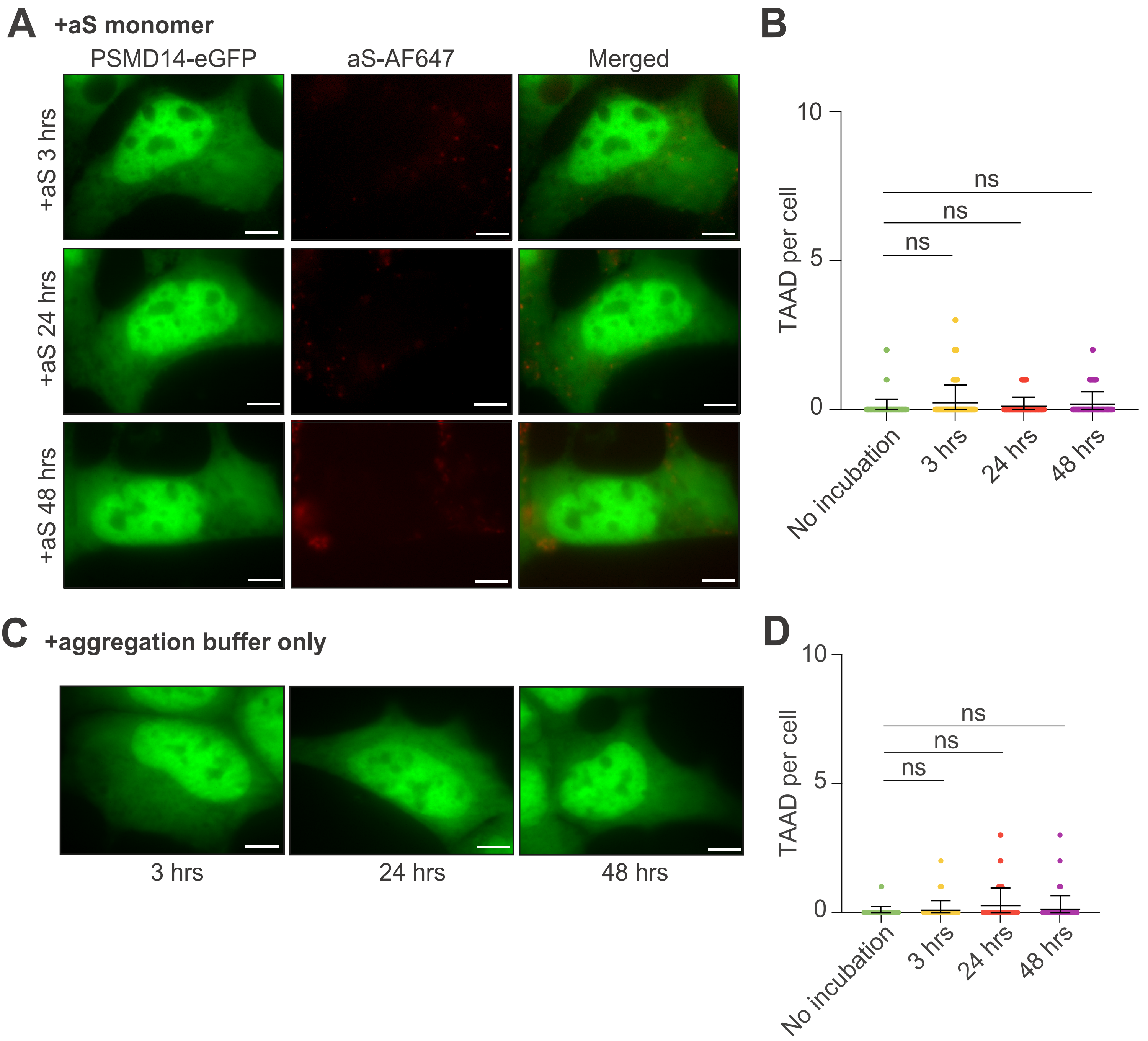

### Supplementary Figure 3

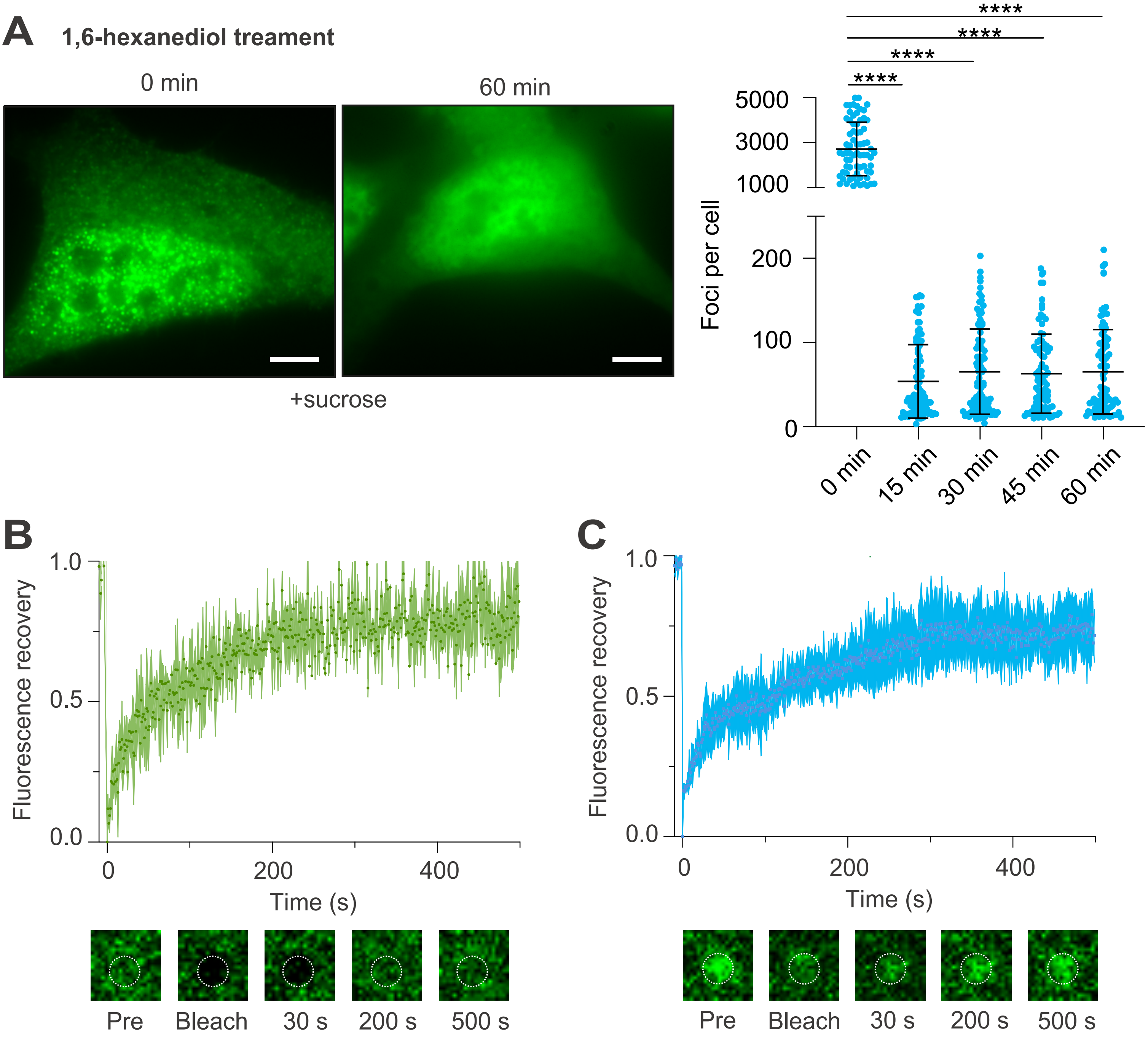

### Supplementary Figure 4

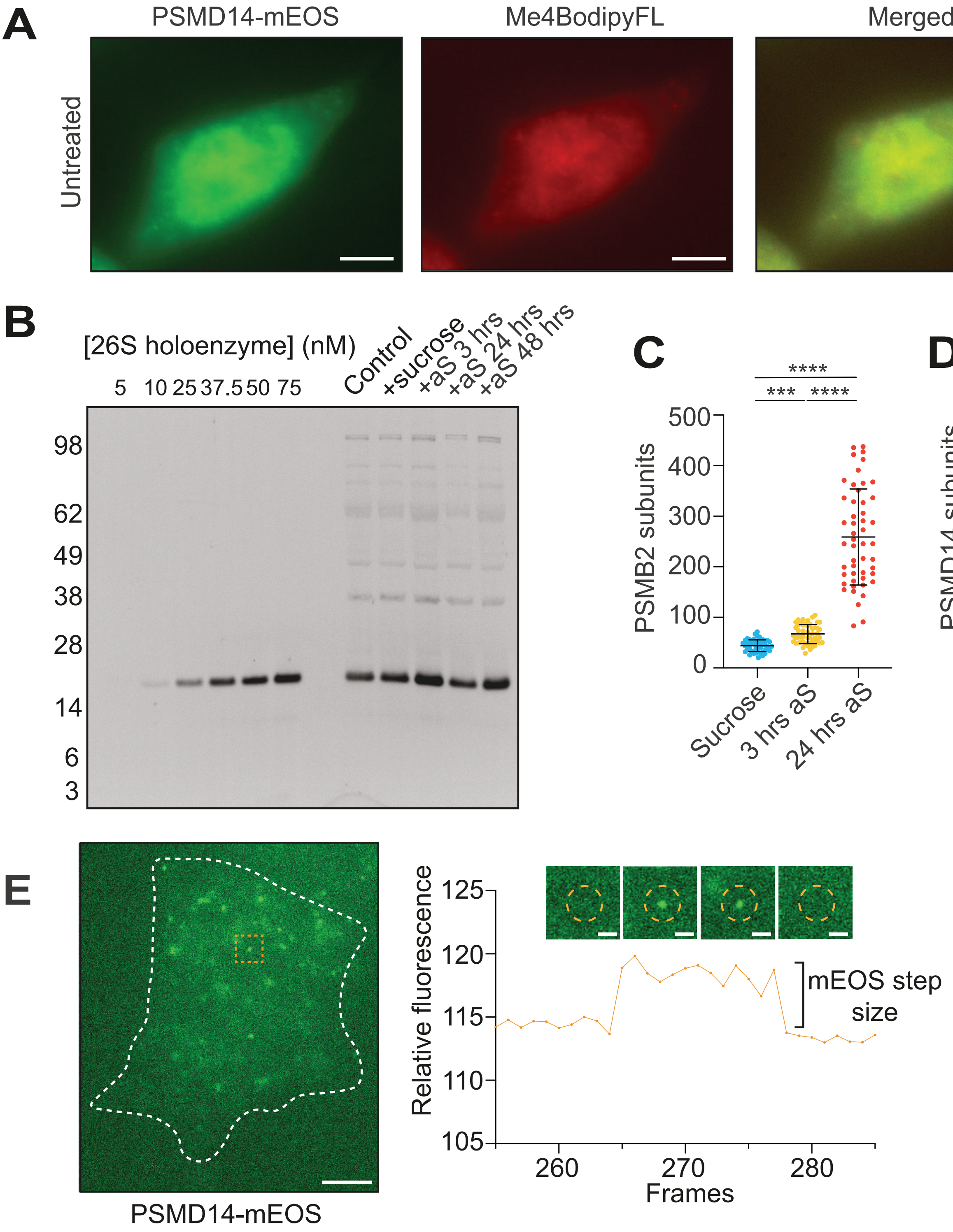

### Supplementary Figure 5

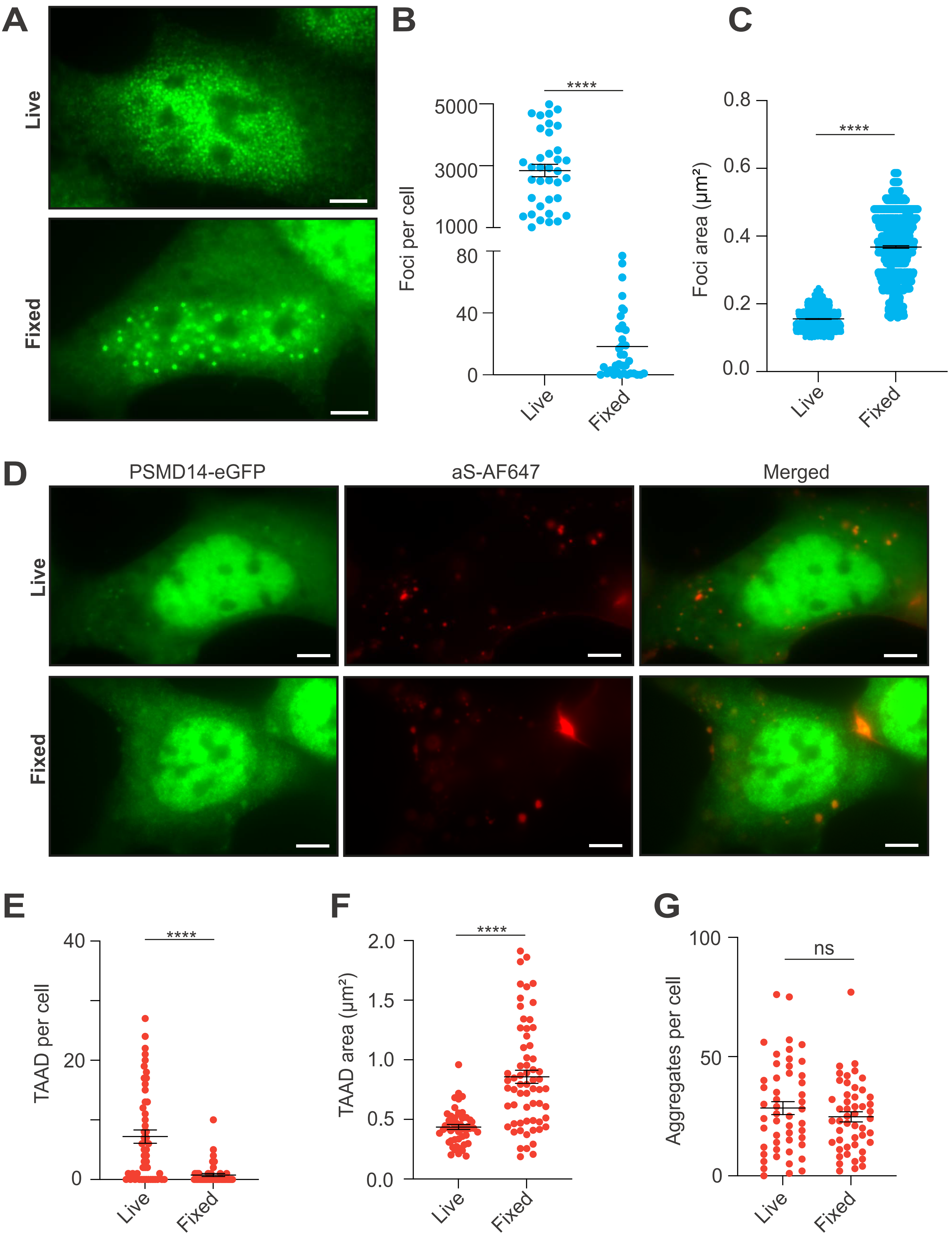

### Supplementary Figure 6

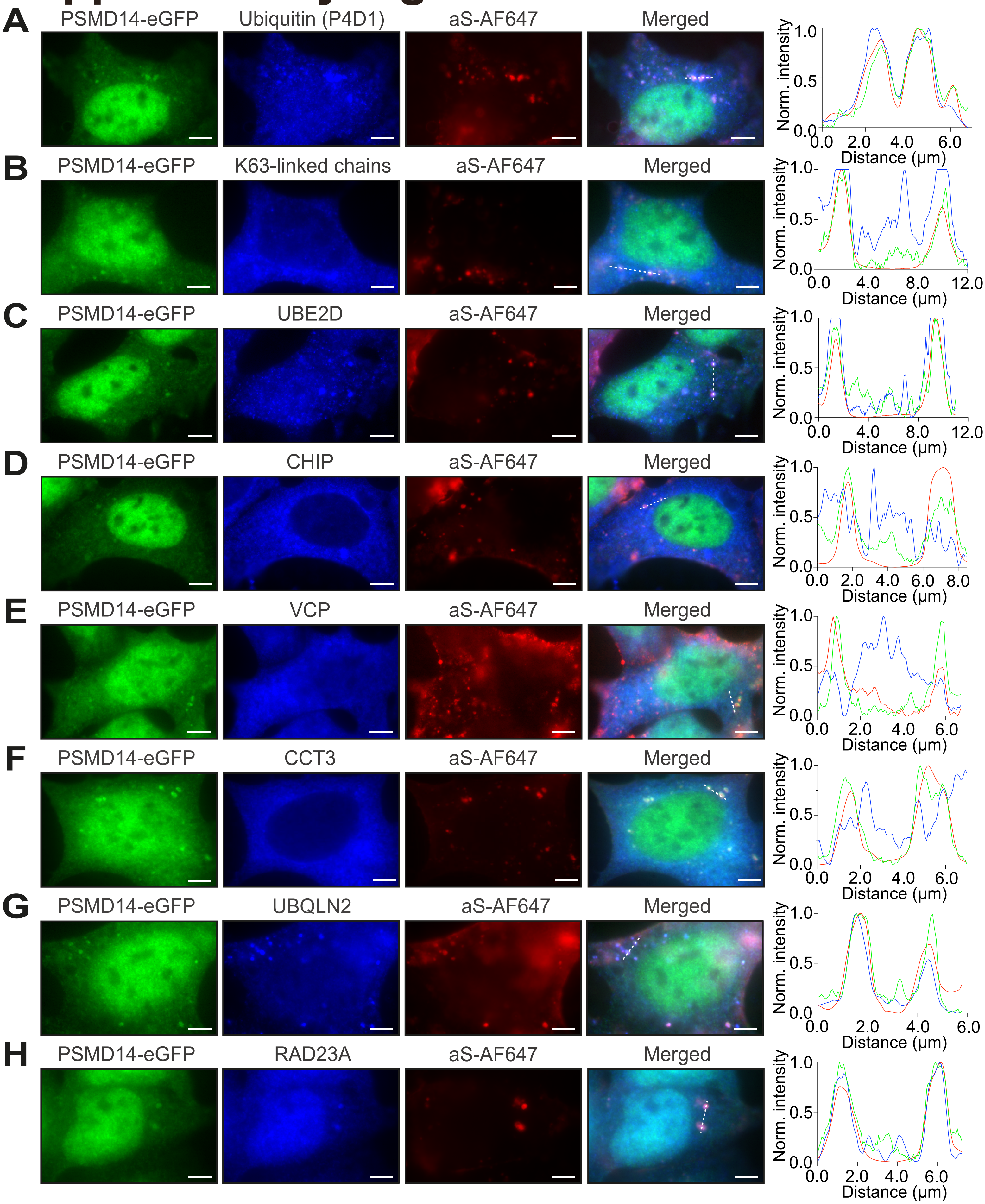

### Supplementary Figure 7

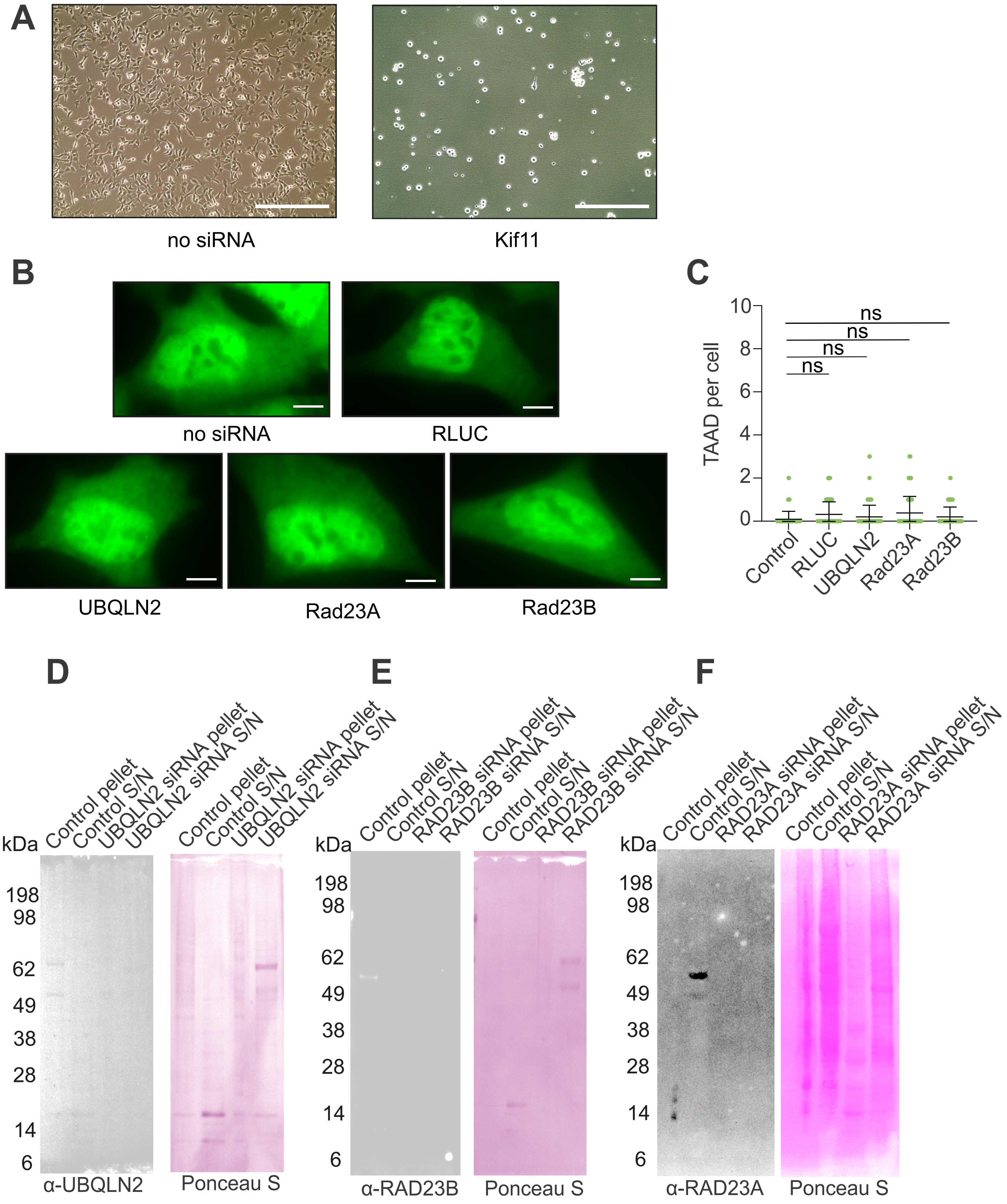

### Supplementary Figure 8

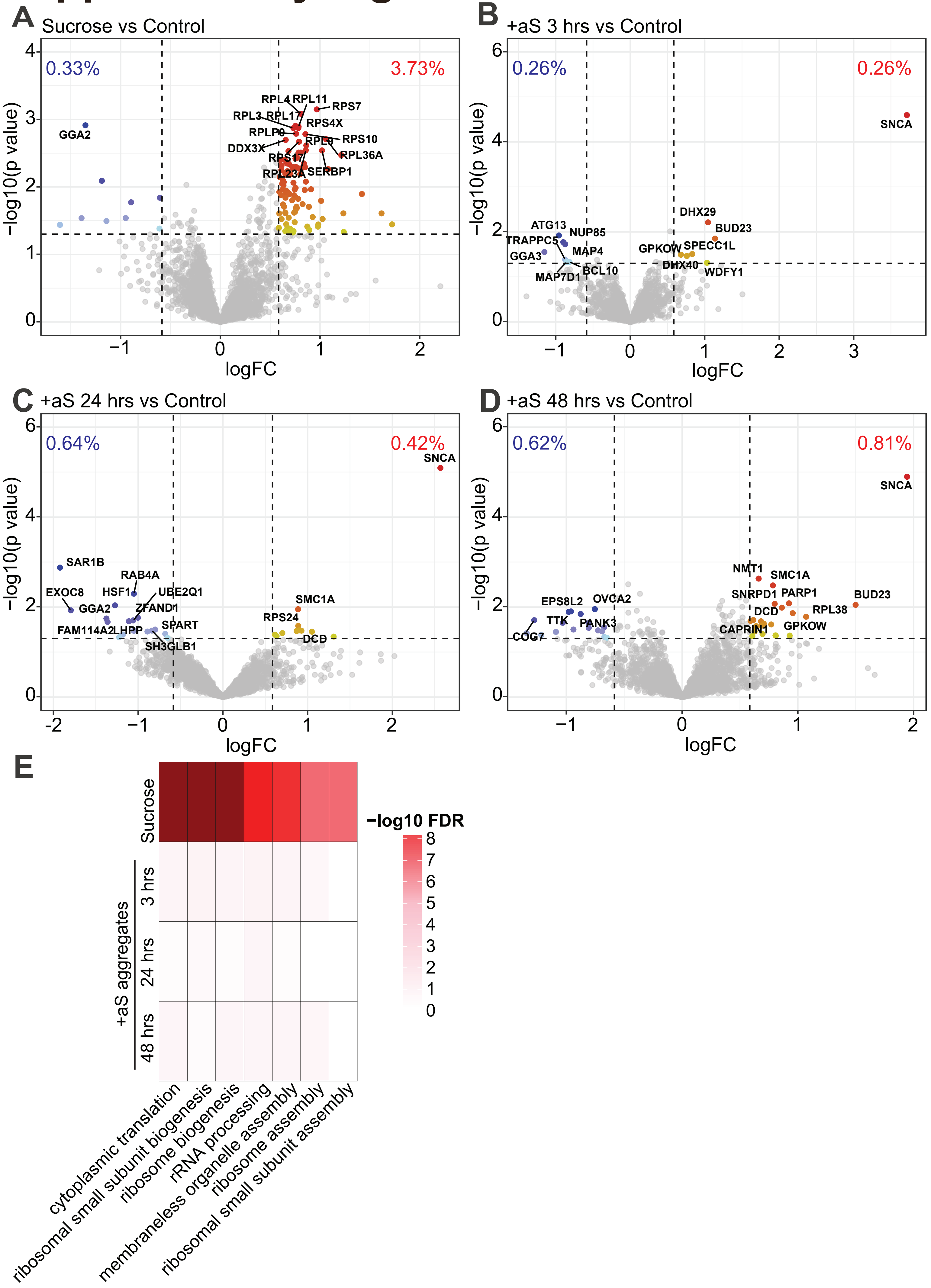

### Supplementary Figure 9

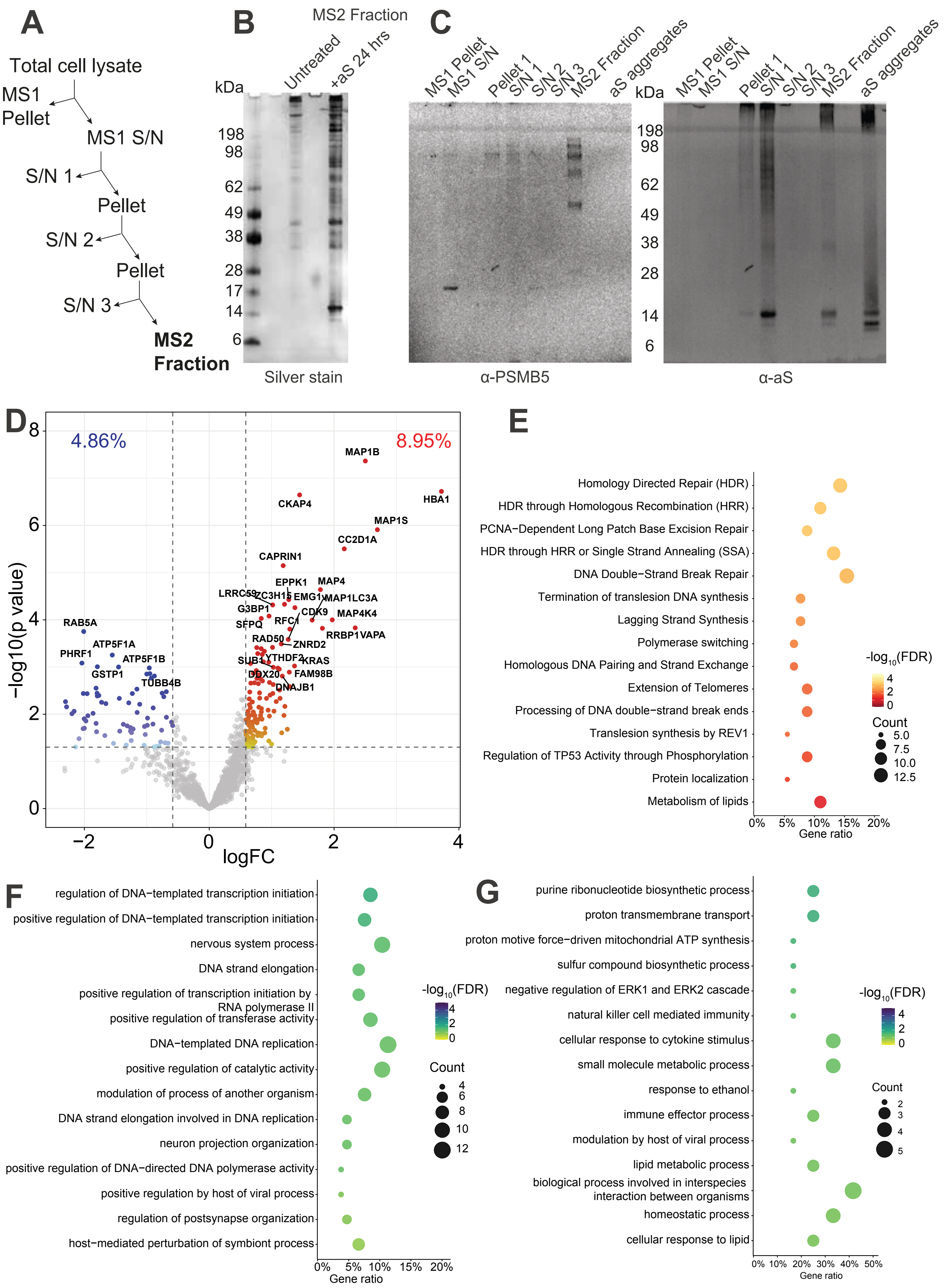

### Supplementary Figure 10

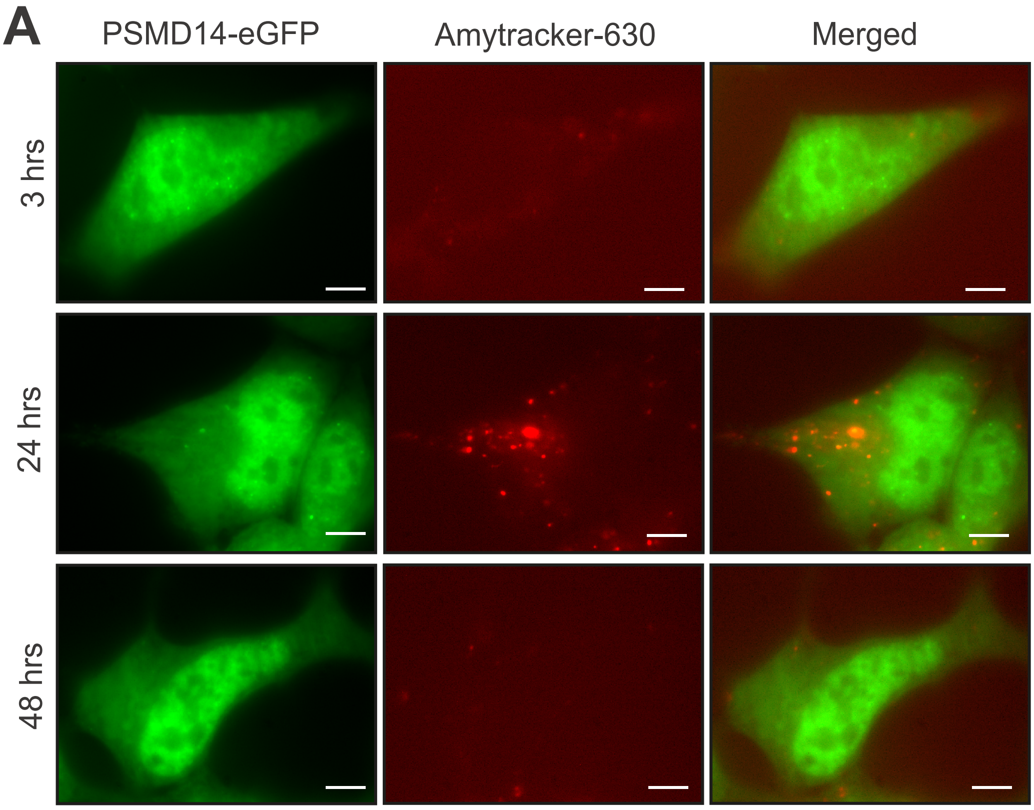


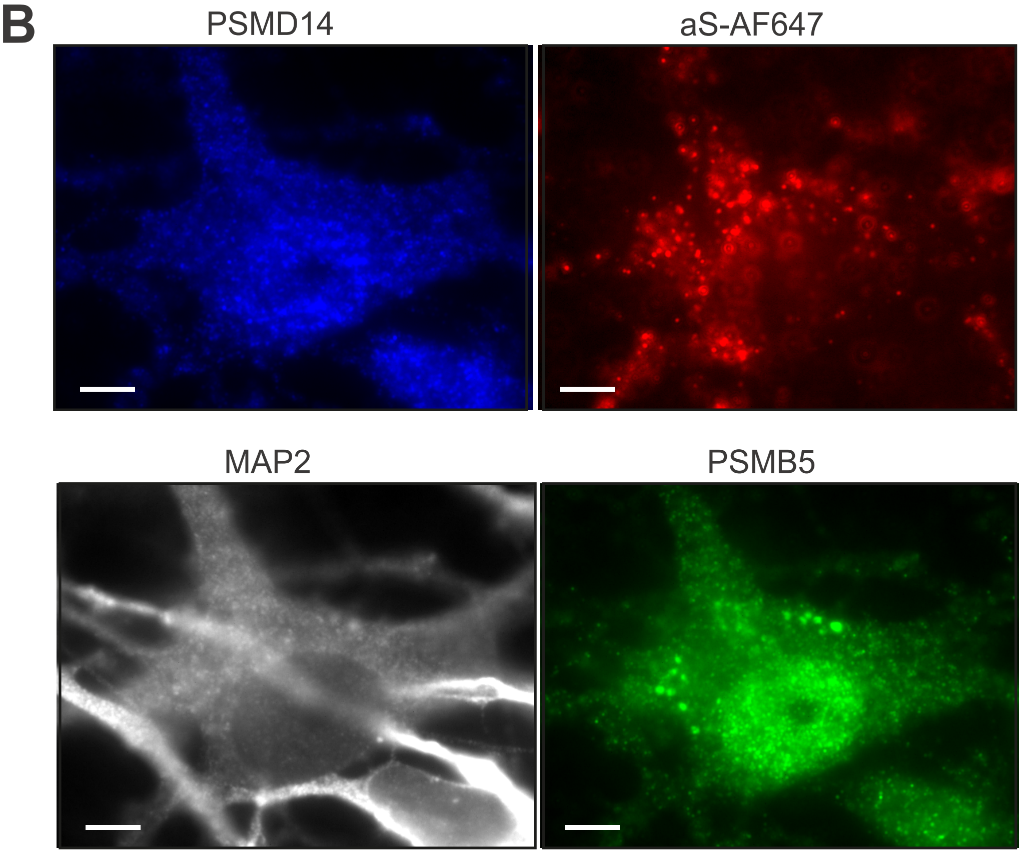

### Supplementary Figure 11

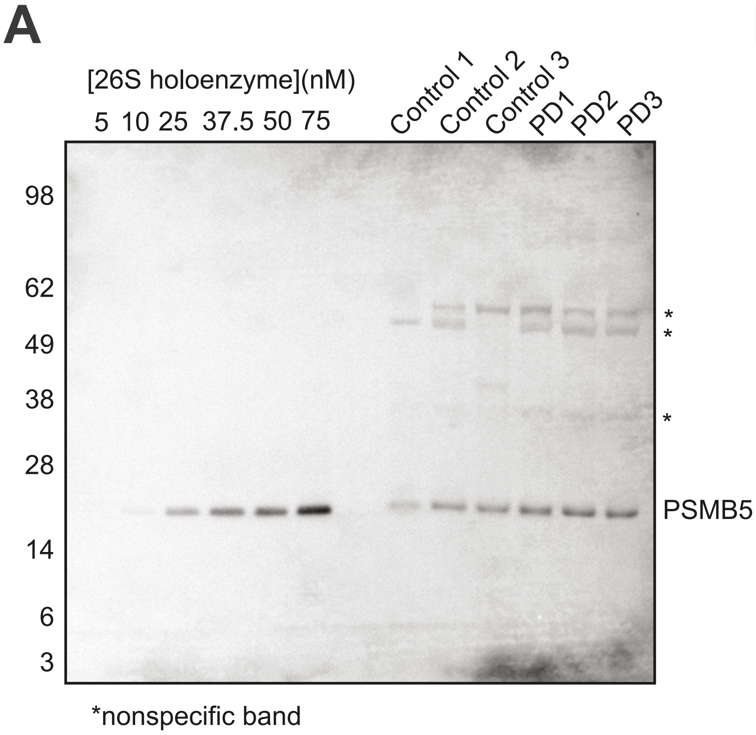


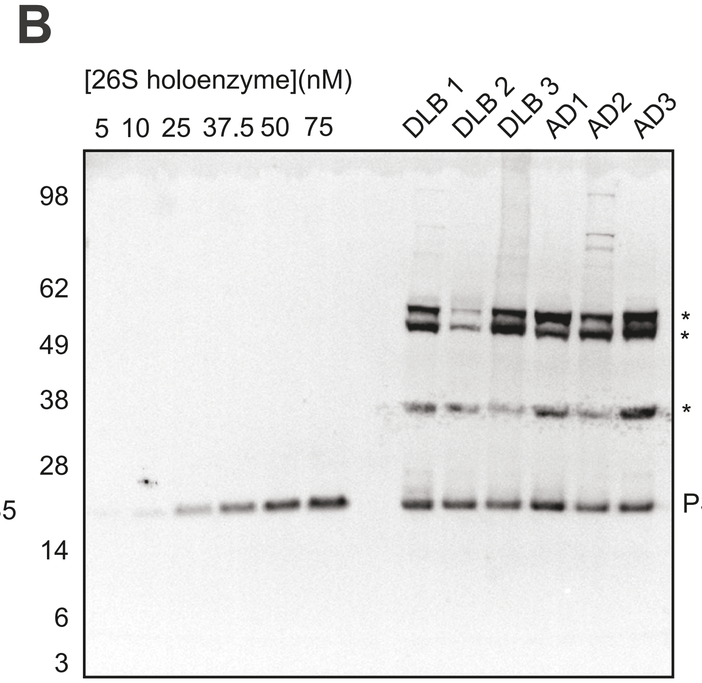
